# Network models predict that pyramidal neuron hyperexcitability and synapse loss in the dlPFC lead to age-related spatial working memory impairment in rhesus monkeys

**DOI:** 10.1101/745901

**Authors:** Sara Ibañez, Jennifer I. Luebke, Wayne Chang, Danel Draguljić, Christina M. Weaver

## Abstract

Behavioral studies have shown spatial working memory impairment with aging in several animal species, including humans. Persistent activity of layer 3 pyramidal dorsolateral prefrontal cortex (dlPFC) neurons during delay periods of working memory tasks is important for encoding memory of the stimulus. *In vitro* studies have shown that these neurons undergo significant age-related structural and functional changes, but the extent to which these changes affect neural mechanisms underlying spatial working memory is not understood fully. Here we confirm previous studies showing impairment on the Delayed Recognition Span Task in the spatial condition (DRSTsp), and increased *in vitro* action potential firing rates (hyperexcitability), across the adult life span of the rhesus monkey. We use a bump attractor model to predict how empirically observed changes in the aging dlPFC affect performance on the Delayed Response Task (DRT), and introduce a model of memory retention in the DRSTsp. Persistent activity—and, in turn, cognitive performance—in both models was affected much more by hyperexcitability of pyramidal neurons than by a loss of synapses. Our DRT simulations predict that additional changes to the network, such as increased firing of inhibitory interneurons, are needed to account for lower firing rates during the DRT with aging reported *in vivo*. Synaptic facilitation was an essential feature of the DRSTsp model, but it did not compensate fully for the effects of the other age-related changes on DRT performance. Modeling pyramidal neuron hyperexcitability and synapse loss simultaneously led to a partial recovery of function in both tasks, with the simulated level of DRSTsp impairment similar to that observed in aging monkeys. This modeling work integrates empirical data across multiple scales, from synapse counts to cognitive testing, to further our understanding of aging in non-human primates.

## 1. INTRODUCTION

Spatial working memory in rhesus monkeys, as in humans, is mediated by the action potential firing activity of neurons in the dorsolateral prefrontal cortex (dlPFC) (reviews: Funahashi, 2017; Constantinidis and Qi, 2018; Miller et al., 2018). And, as in humans (Albert, 1993; Salthouse et al., 2003; Fisk and Sharp, 2004; Rhodes, 2004; Sorel and Pennequin, 2008), the effects of aging on working memory is heterogeneous—while a significant proportion of rhesus monkeys become increasingly impaired on tasks of spatial working memory during normal aging (“unsuccessful agers”), a significant number of “successful agers” show no signs of impairment (Lacreuse et al., 2005; Moore et al., 2006; Moore et al., 2017). Furthermore, for impaired subjects, impairment typically begins fairly early in the aging process, during early middle age (Moore et al., 2005; Moore et al., 2006; Moore et al., 2017). This distribution of impaired and spared monkeys across the lifespan enables assessment of brain changes associated not simply with aging, but with cognitive performance *per se* in this model of normal human aging. This approach has been taken in studies of normal aging by our group and others (reviews: Peters, 2007; Luebke et al., 2010; Wang et al., 2011; Peters and Kemper, 2012; Luebke et al., 2015), in which monkeys were assessed on working memory tasks such as the Delayed Response Task (DRT) and the Delayed Recognition Span Task in the spatial condition (DRSTsp), and their brains subsequently examined for a wide variety of parameters. Thus, declines in spatial working memory have been associated with many sub-lethal changes to the structure and function of neurons, glial cells and white matter pathways in the dlPFC of the rhesus monkey (Peters, 2007; Peters et al., 2008; Peters, 2009; Bowley et al., 2010; Luebke et al., 2010; Shobin et al., 2017). Reductions in synapses and increased dystrophy of white matter pathways begin in early middle age; for example, Peters et al. (Peters et al., 2008) showed a continuous decrease in the number of excitatory and inhibitory synapses, detectable even in middle age in the monkey dlPFC.

In addition to well-documented structural changes, functional alterations to the electrophysiological properties of supragranular neurons in the aging monkey dlPFC have been reported in several studies (Chang et al., 2005; Wang et al., 2011; Coskren et al., 2015). Layer 3 dlPFC pyramidal neurons exhibit significantly increased evoked action potential (AP) firing rates *in vitro* for aged compared to young monkeys (Chang et al., 2005; Coskren et al., 2015). In contrast, another study (Wang et al., 2011) reported that *in vivo* firing rates of dlPFC DELAY neurons decreased with aging during the DRT, for monkeys performing the task well. Since persistent firing patterns of dlPFC pyramidal neurons encode precisely tuned spatial and temporal information during a working memory task, exploring how these findings might complement each other may reveal new predictions about spatial working memory decline.

While the growing body of literature over the past 20 years has documented numerous changes to neurons, glial cells and white matter pathways in the aging brain that are associated with cognitive changes, which changes are the key determinants of age-related cognitive declines has not yet been firmly established (Hof and Morrison, 2004; Luebke et al., 2010; Morrison and Baxter, 2012; Peters and Kemper, 2012; Konar et al., 2016; Motley et al., 2018; Cleeland et al., 2019). This lack of insight is partially due to the difficulty in first, selectively targeting individual variables (such as firing rate or synapse number) in humans or experimental subjects without, second, also altering upstream or downstream effectors. A powerful way to gain insight into which age-related changes are most consequential for dlPFC network behavior during normal aging is through computational models that are constrained by empirical data. This is the approach we use here through a systematic study of the “bump attractor” neural network model, and its extension to memory retention in a new task similar to the DRSTsp.

The bump attractor network has been used to model spatial working memory in the DRT (Compte et al., 2000; Wang et al., 2011; Wang et al., 2013; Wimmer et al., 2014; Wu et al., 2016 for review). The bump attractor exhibits persistent firing activity after an initial stimulus disappears, encoding a memory of the stimulus location. The bump attractor was used to show that an age-related loss of synaptic strength could account for reduced firing rates of dlPFC DELAY neurons (Wang et al., 2011), but the effects of increased excitability of pyramidal neurons seen *in vitro* on the function of bump attractor models has not been examined. We have previously used computational modeling to predict which intrinsic electrophysiological and morphological properties of individual pyramidal neurons contribute to the action potential firing rate increases seen in aging (Coskren et al., 2015; Rumbell et al., 2016). Here we extend these previous single neuron studies using network-level modeling to predict the effect of empirically observed physiological and structural changes in the aging rhesus dlPFC on cognitive behavior. We obtained whole-cell patch clamp recordings of dlPFC pyramidal neurons from behaviorally characterized monkeys across the adult lifespan. Empirical results indicated that previously reported physiological changes seen in aging are already present in middle age, and are correlated with cognitive impairment. This modeling work aims to connect age-related changes in non-human primates across multiple empirical scales, from synapse counts and physiology of single neurons up to network output and cognitive performance. We predict that the increased firing rates and reduced synapse density observed with aging may partially compensate one another, but still are sufficient to induce substantial cognitive deficits in the DRT and DRSTsp.

## 2. MATERIALS AND METHODS

### 2.1 Experimental Subjects

The rhesus monkeys (*Macaca mulatta*) studied here were a part of a larger study of normal aging of the brain. Data were obtained from a total of 9 young (8.9 ± 0.5 years old), 18 middle-aged (18.2 ± 0.5 years old), and 12 aged (24.7 ± 0.7 years old) monkeys (Table 1). Prior to the electrophysiological experiments, all but 3 young monkeys completed cognitive testing of the Delayed Recognition Span Task in the spatial condition (DRSTsp; Moore et al., 2017). Monkeys were obtained from the Yerkes National Primate Research Center at Emory University (Atlanta, GA, USA), and housed in the Laboratory Animal Science Center (LASC) at the Boston University School of Medicine (BUSM) under strict accordance with the guidelines established by the NIH *Guide for the Care and Use of Laboratory Animals* and the US *Public Health Service Policy on Humane Car and Use of Laboratory Animals*. Both Yerkes National Primate Research Center and BUSM LASC are fully accredited by the Association for Assessment and Accreditation of Laboratory Animal Care.

**Table 1.** Experimental Subjects

|  | Monkey | Age (y) | Sex | DRST Spatial |
| --- | --- | --- | --- | --- |
| Young: | PIL | 6.8 | M | - |
|  | PIE | 6.9 | M | - |
|  | PIK | 8.2 | M | - |
|  | AM299 | 8.6 | F | 2.50 |
|  | AM296 | 8.8 | M | 2.22 |
|  | AM289 | 9 | M | 3.05 |
|  | AM255 | 9.5 | F | 3.42 |
|  | AM295 | 10.9 | M | 2.27 |
|  | AM254 | 11.4 | F | 2.40 |
| Middle Aged: | AM321c | 13.4 | M | 3.61 |
|  | AM278 | 15.3 | F | 2.31 |
|  | AM288 | 16 | M | 2.22 |
|  | AM351c | 16.4 | F | 1.94 |
|  | AM285 | 17 | F | 2.14 |
|  | AM350c | 17.3 | F | 2.92 |
|  | AM340 | 17.6 | F | 2.12 |
|  | AM297 | 18.1 | M | 2.26 |
|  | AM265L | 18.2 | F | 1.84 |
|  | AM272L | 18.2 | F | 2.32 |
|  | AM274 | 19 | M | 2.31 |
|  | AM342c | 19.2 | F | 2.94 |
|  | AM279 | 19.5 | M | 2.43 |
|  | AM271L | 20 | F | 1.95 |
|  | AM263L | 20.3 | F | 2.02 |
|  | AM311c | 20.3 | M | 3.19 |
|  | AM314x | 20.8 | M | 2.71 |
|  | AM264L | 20.9 | F | 2.07 |
| Aged: | AM270L | 21 | F | 2.06 |
|  | AM266L | 21.1 | F | 2.06 |
|  | AM286 | 22.4 | F | 2.1 |
|  | AM282 | 23.1 | M | 2.20 |
|  | AM281 | 23.2 | M | 2.23 |
|  | AM276 | 24.4 | M | 2.12 |
|  | AM356 | 24.8 | M | 1.99 |
|  | AM284 | 25.5 | M | 2.28 |
|  | AM283 | 26.4 | M | 2.04 |
|  | AM 298 | 26.8 | M | 2.38 |
|  | AM355 | 28.4 | M | 2.08 |
|  | AM034L | 29.5 | M | 2.18 |

### 2.2 Behavioral Assessment

A total of 36 subjects were tested on the DRSTsp, 6 young, 18 middle-aged, and 12 aged monkeys. The DRSTsp is a short-term memory task in which the subject distinguishes a novel cue (spatial location) from an increasing set of recently presented, familiar cues (spatial locations) (Moore et al., 2017 and Fig. 1a). All stimuli were identical brown disks, each covering one of 18 small wells arranged on a testing board in a 3×6 matrix. On the first trial one of the wells was baited with a reward then covered with a disk. Once a screen was raised, the monkey moved the disk to receive the reward. The screen was lowered before the second trial, when that same well (with no reward) was covered with a disk and a second well was baited with a reward and covered with an identical disk (10 s delay). Subsequent trials include the previous cue plus one new cue, until the monkey is unable to identify novel cues (spatial locations). Monkeys were tested for 10 consecutive days. The cognitive score recorded is the recognition span: the mean number of unique cues the monkey identified correctly before making a mistake. Some of the subjects were assessed on the DRSTsp more than once over a period of several years as part of a longitudinal study of aging. The DRSTsp score used in the present study for longitudinally assessed subjects was the one most proximal to the date the subject was sacrificed.

**Figure 1:**
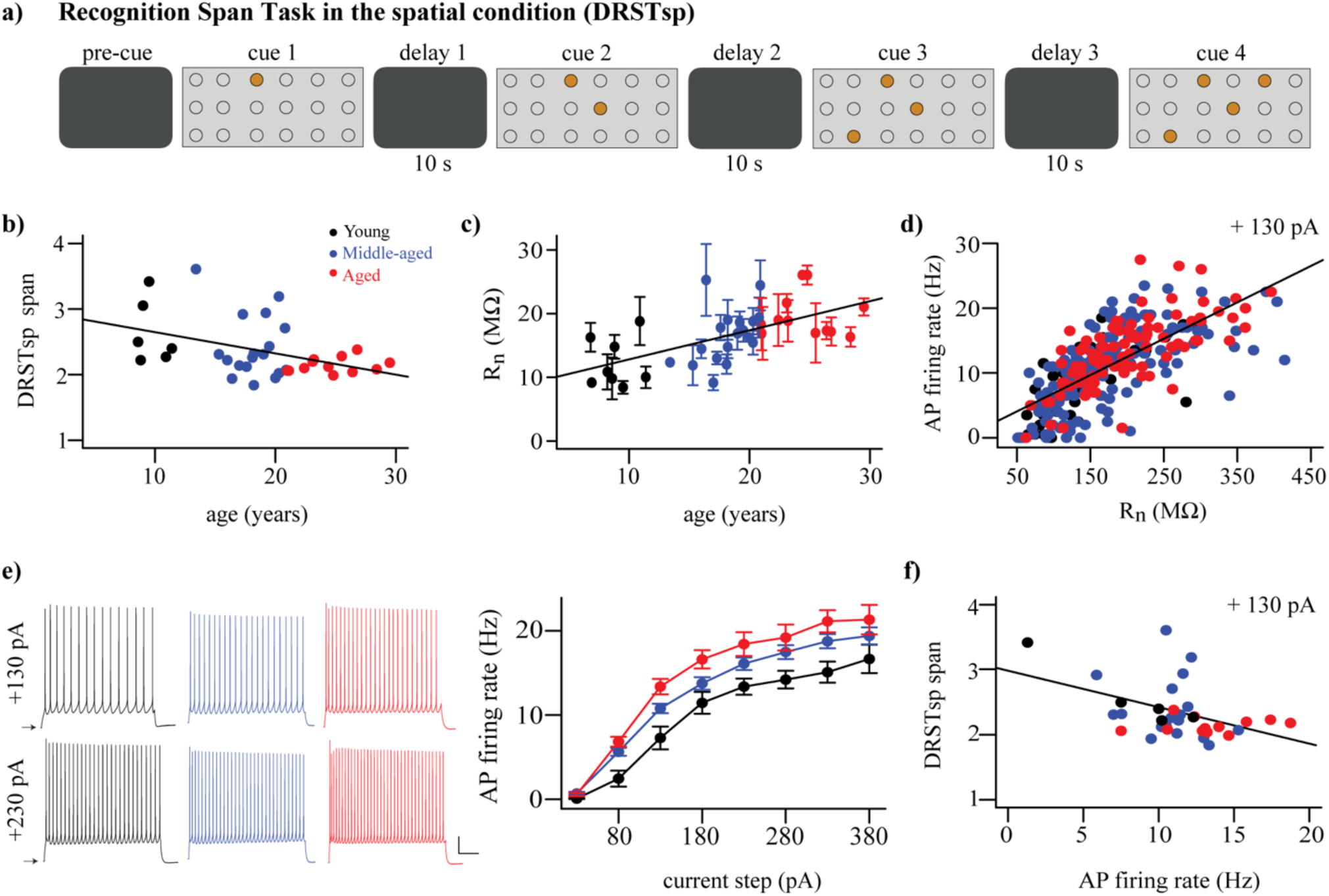
Cognitive performance and physiological variables vary with aging. (a) DRSTsp setup. Brown circles represent disks covering the wells arranged on a board in a 3×6 matrix. (b) DRSTsp performance vs. age. (c) Input resistance vs. aging; regression includes subject as random effect. Data shown as mean ± SEM for each subject. (d) FR vs. input resistance for + 130 pA current step; these variables were highly correlated at all levels of current injection (not shown). (e) Left: Trains of APs evoked by 2-s depolarizing current steps for representative young, middle-aged and aged neurons. Scale bar: 50 mv and 500 ms. Current steps were applied from resting membrane potentials, shown as arrows. Right: FR vs. injected current. (f) DRSTsp vs. AP firing rate in response to a +130 pA current injection. In all graphs, young, middle-aged, and aged subjects are shown in black, blue, and red respectively. Data were obtained from a total of 9 young, 18 middle-aged, and 12 aged monkeys (Table 1). All but 3 young monkeys completed cognitive testing of the Delayed Recognition Span Task in the spatial condition. Whole-cell patch clamp recordings from 324 pyramidal neurons from 38 subjects were used.

### 2.3 Preparation of Cortical Slices and Whole-cell Patch Clamp Experiments

Monkeys were sacrificed as described in our previous publications (Amatrudo et al., 2012; Medalla and Luebke, 2015; Medalla et al., 2017). Briefly, following perfusion with Krebs-Henseleit solution, a block of tissue (∼1cm^3^) was removed from the left lateral prefrontal cortex, and sectioned in ice-cold oxygenated Ringer’s solution (concentrations, in mM: 26 NaHCO_3_, 124 NaCl, 2 KCl, 3 KH_2_PO_4_, 10 glucose, 1.3 MgCl_2_, pH 7.4; Sigma-Aldrich) into 300µm-thick coronal slices with a vibrating microtome. Slices were immediately transferred into room temperature, oxygenated Ringer’s solution, for a minimum of one hour equilibration prior to recordings.

Individual slices were placed into submersion-type recording chambers (Warner Instruments) positioned on Nikon E600 infrared-differential interference contrast (IR-DIC) microscopes. During recordings, slices were continuously superfused with oxygenated, room-temperature Ringer’s solution at a flow rate of 2-2.5 ml/min.

Standard tight-seal, whole-cell patch clamp recordings were performed on layer 3 pyramidal neurons as described previously (Amatrudo et al., 2012; Luebke et al., 2015; Medalla and Luebke, 2015; Medalla et al., 2017). Electrodes were fabricated using a Flaming/Brown micropipette puller (MODEL P87, Sutter Instruments), and filled with potassium methane sulfonate (KMS)-based internal solution (concentration, mM: 122 KCH_3_SO_3_, 2 MgCl_2_, 5 EGTA, 10 NaHEPES, 1% biocytin, pH 7.4; Sigma-Aldrich). Experiments were performed using EPC-9 or EPC-10 amplifiers (HEKA Elektronik) controlled with PatchMaster acquisition software (HEKA Elektronik). Signals were low pass-filtered at 10kHz, and access resistance was monitored and maintained throughout each experiment. Current clamp protocols were implemented as previously described (Amatrudo et al., 2012; Luebke et al., 2015; Medalla and Luebke, 2015; Medalla et al., 2017), to assess the input resistance (in MΩ) and repetitive firing rate (FR, measured in Hz) of the neurons.

Whole-cell patch clamp recordings from 324 layer 3 dlPFC pyramidal neurons from 38 subjects were used in this study. In total, 40 neurons for the young, 188 for the middle-aged and 96 for the aged subjects.

### 2.4 The bump attractor network model

The “bump attractor” network model was originally designed to simulate a spatial working memory task for awake behaving monkeys known as the Delayed Response Task (DRT) or the Oculomotor Delay Task (Fig. 2a, Goldman-Rakic, 1995; Compte et al., 2000; Wang et al., 2011; Wimmer et al., 2014). After the monkey fixates on the center of a computer screen (for 1 s in our simulations), a light appears during the cue period in one of eight directions around a circle (occurring at 0° and lasting 0.5 s in our simulations). After the cue period the light turns off, and the monkey fixates on the center of the screen during a delay period (2 s in our simulations). During the response period (0.7 s in our simulations), the monkey shifts its gaze (makes a saccade) to indicate its memory of the location of the stimulus. The selective firing of pyramidal neurons in the dlPFC to different spatial locations (their preferred directions) is thought to be the mechanism by which stimulus location is encoded. Thus, monkeys performing this task successfully have neurons that exhibit persistent neural activity during the delay period tuned to the stimulus location (Goldman-Rakic, 1995; Compte et al., 2000; Wimmer et al., 2014). This is achieved in the bump attractor model with excitatory neurons tuned to each spatial location (so that we can label each neuron by that location— its “preferred direction”), with the synaptic strength between pairs of excitatory neurons determined by the difference between the neurons’ preferred directions.

**Figure 2:**
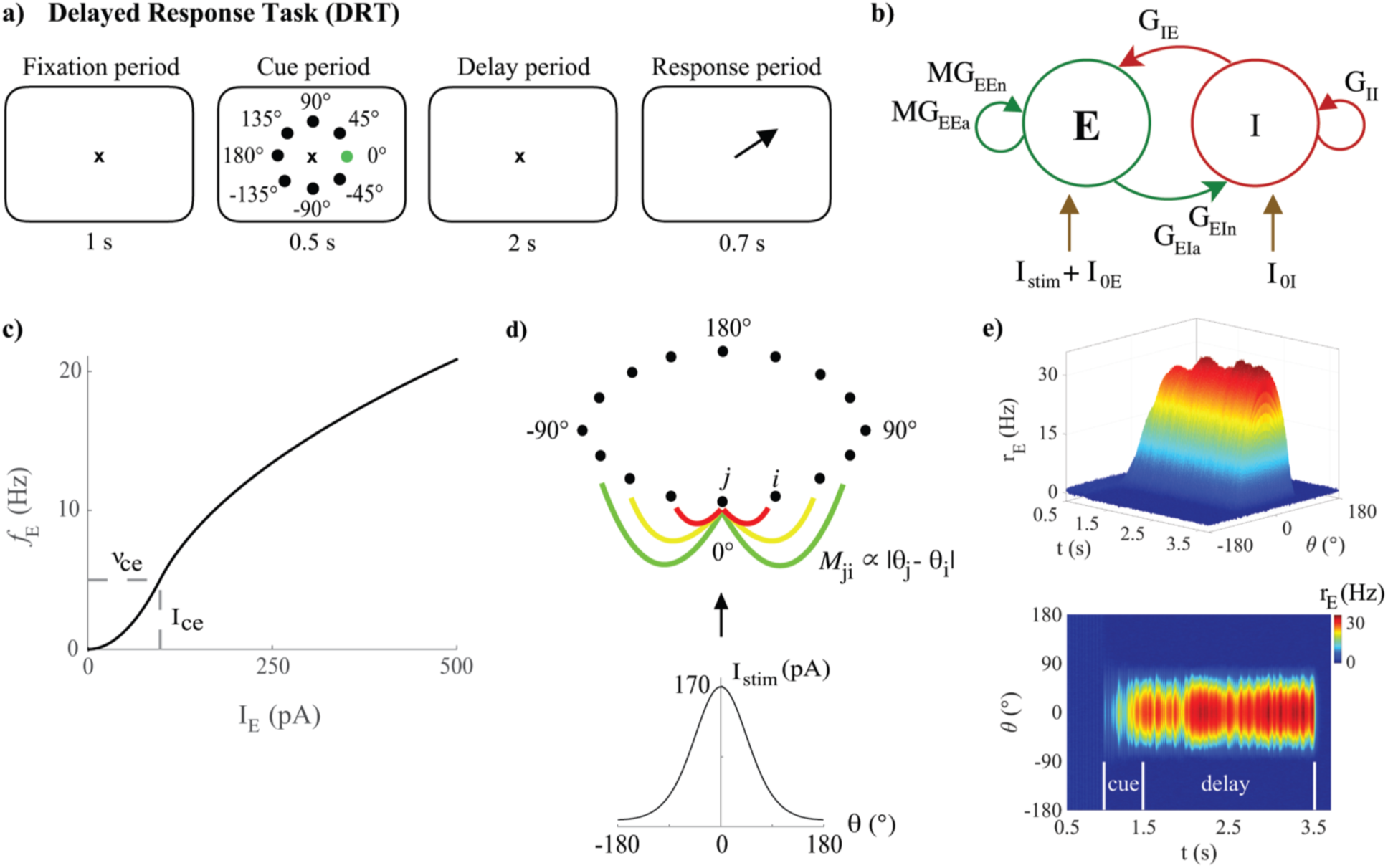
The bump attractor model of the DRT. (a) The DRT includes fixation, cue, delay and response periods of 1 s, 0.5 s, 2 s, and 0.7 s respectively. The ×, black dots, and green dot respectively represent the fixation point, the 8 possible stimulus locations, and actual stimulus location (0°) in the simulations. Arrow indicates the saccade direction during the response period. (b) The recurrent excitatory-inhibitory network. (c) The *f*-I curve for young model excitatory neurons. (d) Ring connectivity among excitatory neurons. *M*_*ij*_ is the connectivity matrix element between excitatory neurons *j* and *i*, depending on the difference between their preferred directions (*θ*_*j*_ and *θ*_*i*_, respectively). Each neuron is strongly connected to its nearest neighbors (red line), with the connection strength decaying with distance (yellow and green lines). The neuron with preferred direction 0° receives the strongest stimulus current, which follows a Gaussian distribution centered at 0°, generating a bump of activity during the delay period (persistent activity tuned to the stimulus location) when excitation and inhibition in the network are well-balanced. € Sample DRT model output (side and top views) when the 0° stimulus location is encoded correctly: excitatory neuron FR vs. simulation time with neurons labeled by their preferred direction.

The bump attractor network used here assumed a firing-rate model for individual neuronal dynamics, modified from (Wimmer et al., 2014). The code was written in Matlab and is available on ModelDB (McDougal et al., 2017) at Accession #256610. The local cortical network was composed of N_E_ = 640 excitatory (pyramidal) neurons (80%) and N_I_ = 160 inhibitory neurons (interneurons, 20%) (Abeles, 1991; Braitenberg and Schütz, 1991) (Fig. 2b). Each excitatory and inhibitory neuron received three general types of synaptic inputs: from excitatory and inhibitory connections with other neurons in the same network, and excitatory connections from other unspecified areas external to the network. The modeled excitatory neurons also received an external input current representative of the “cue”. The synaptic excitation of each neuron was modeled with two distinct equations, separately representing synaptic inputs mediated by AMPA and NMDA receptors. The voltage dependence of the NMDA currents was omitted from the present study, since Compte et al. (Compte et al., 2000) showed that the slow kinetics of NMDA-mediated synaptic transmission was the most important feature for generating stable persistent activity.

The equations for FR of each excitatory and inhibitory neuron (r_E_ and r_I_) were given by

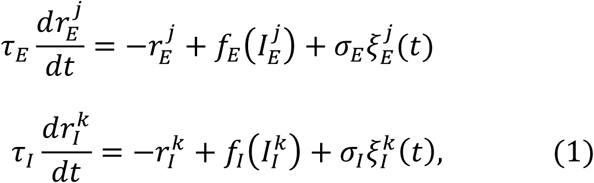

for j = 1, …, N_E_ and k = 1, …, N_I_. The parameters *τ*_*E*_ and *τ*_*I*_ represented the membrane time constants for excitatory and inhibitory neurons respectively (in ms), with *f*(*I*) the FR activation function (or *f-*I curve, in Hz), total synaptic input currents I_E_ and I_I_ (in pA), and Gaussian white noise inputs 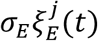 and 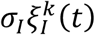 (in Hz), where *σ*_*E*_ and *σ*_*I*_ were the standard deviations. In this study, *τ*_*E*_ = 20 ms, *τ*_*I*_ = 10 ms, *σ*_*E*_ = 1, and *σ*_*I*_ = 3. The activation function for the firing rate of each neuron took the piecewise form (Brunel, 2003, Wimmer et al., 2014, Roxin and Compte, 2016):

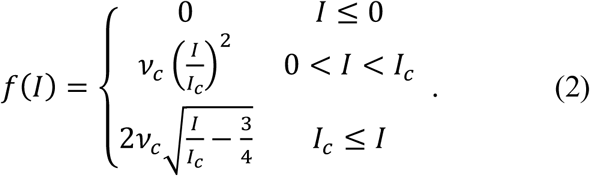

The parameter I_c_ represented a current near rheobase (measured in pA), with corresponding firing rate *ν*_c_ (in Hz) (see Fig. 2c). Thus, the parameter pairs {*ν*_ce_, I_ce_} and {*v*_ci_, I_ci_} uniquely defined the *f-*I curves for excitatory and inhibitory neurons. Here, I_ci_ = 20 pA and *v*_ci_ = 50 Hz, values approximately in the regime for Chandelier cells in the monkey PFC (Zaitsev et al., 2009, Povysheva et al., 2013). The parameters *ν*_ce_ and I_ce_ were chosen to fit our empirical data as described below. See also the supplementary information for a general, less excitable form of both *f*-I curves.

The total synaptic input currents (in pA) for each neuron were

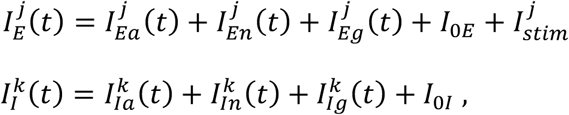

with time-dependent synaptic currents given by

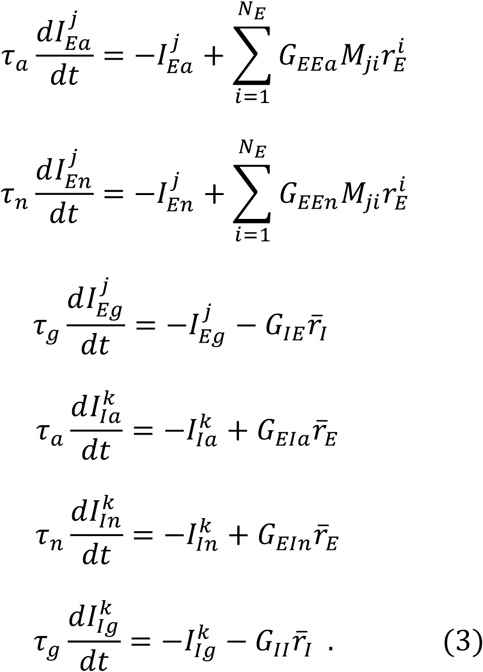

The subindices *a* and *n* represented the AMPA and NMDA synaptic receptors for the glutamatergic synapses, and *g* the GABA_A_ receptor. Parameters *τ*_*a*_, *τ*_*n*_ and *τ*_*g*_ were the synaptic decay time constants for AMPA, NMDA and GABA_A_ (Hestrin et al., 1990; Spruston et al., 1995; Salin and Prince, 1996; Xiang et al., 1998). We set *τ*_*a*_ = 2 ms, *τ*_*n*_ = 100 ms, *τ*_*g*_ = 10 ms. The quantity 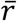 represented the mean FR value of each neuron type, and the *G*_*pq*_ parameters with {p, q} = {E, I} were the constant synaptic weights. The connectivity matrix *M* was a circular Gaussian function describing the translation-invariant connections among the excitatory neurons. Labeling each neuron by its preferred direction, the matrix element *M*_*ji*_ for two neurons *j* and *i* depended on the difference between their preferred directions. That is, *M*_*ji*_ ∝ |*θ*_*j*_ – *θ*_*i*_|, with

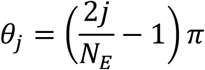

being the preferred stimulus direction of the excitatory neuron *j*, where a complete circumference was divided into N_E_ angles (see Fig. 2d). The quantities I_0E_ and I_0I_ represented constant excitatory input currents from other brain areas to excitatory and inhibitory neurons respectively, and 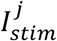 the input current to each excitatory neuron due to stimuli during cue periods of the tasks described below. Here, I_0E_ = 80 pA and I_0I_ = 15 pA. The parameters *G*_*pq*_ were varied using the Latin Hypercube Sampling design, as described below.

The input stimulus currents for each excitatory neuron representing a stimulus at 0° was modeled as in (Wimmer et al., 2014) as

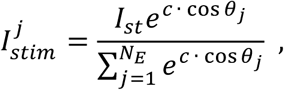

which provided the bump-shaped input current to all excitatory neurons (Fig. 2d). Parameter I_st_ defined the strength of the stimulus and *c* the concentration of excitatory neuron to excitatory neuron connectivity. In our simulations, I_st_ = 40000 pA, and *c* = 1.5 for the DRT model and *c* = 20 for the DRSTsp model. A typical simulation of the DRT is shown in Figure 2e.

### 2.5 Modeling effects of aging on individual neurons

#### Increased AP firing rate (hyperexcitability) of pyramidal neurons with aging

We fit parameters *ν*_ce_ and I_ce_ of the excitatory neurons *f-*I curve to *in vitro* FR data from the subjects described above. We computed the mean FR vs. different input currents of all neurons from each monkey, then computed a grand mean of FR vs. input current for monkeys in each age group (Fig. 3b). Fits of the parameters *ν*_ce_ and I_ce_ to these data were estimated manually using the FindFit function from Mathematica (Wolfram Research, Champaign, IL). The value I_ce_ = 98 pA fit all three age groups well, and also agreed with rheobase values for regular spiking pyramidal neurons with low input resistance (Zaitsev et al., 2012). We used *ν*_ce_ = 5, 7, and 9 Hz to fit the *f-*I curve to data from the young, middle-aged, and aged groups respectively (Fig. 3b). Thus, to fit *in vitro* FR data it was sufficient to increase the single parameter *ν*_ce_. Assuming the magnitude of changes to *f-*I curves with aging would be similar at physiological vs. room temperature, we used these parameter values directly in our ‘young’, ‘middle-aged’ and ‘aged’ network models. We also repeated some DRT simulations after fitting a generalized, less excitable firing rate activation function to the young, middle-aged, and aged monkey data (supplementary information and supplemental Fig. 1a).

**Figure 3:**
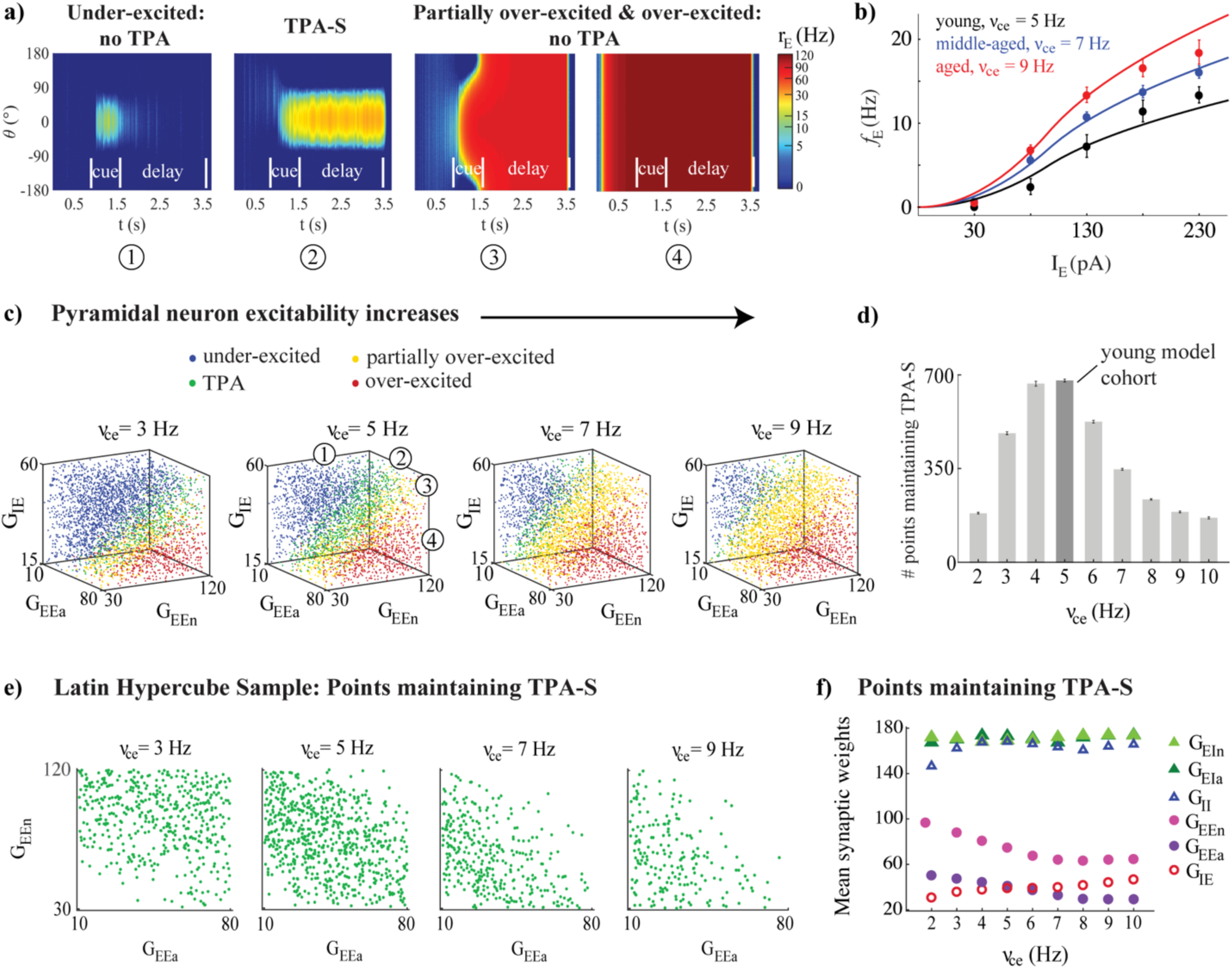
DRT model output varied across the parameter space. (a) From left to right: Example model output showing (1) under-excitation; (2) TPA-S; (3) partial over-excitation; and (4) over-excitation. (b) The model *f*-I curve fit (solid lines) to empirical AP firing rates of pyramidal neurons of young, middle-aged and aged subjects, averaged for each age group (black, blue, and red respectively, shown as mean ± SEM at each injection level). (c) DRT model output for 4200 points of the parameter space LHS, shown in 3D projections across the subspace of the excitatory (G_EEa_ and G_EEn_) and the inhibitory (G_IE_) synaptic weights of pyramidal neurons as their excitability increased (*ν*_ce_ = 3, 5, 7, and 9 Hz). The numbers in the 3D projection for *ν*_ce_ = 5 Hz indicate each type of network performance shown in (a), represented as blue, green, yellow, and red dots respectively in each graph. (d) Number of points in each LHS maintaining TPA-S as *ν*_ce_ increased. Dark gray bar indicates the young model cohort. (e) Projections across the (G_EEa_, G_EEn_) subspace showed that points maintaining TPA-S shifted to lower (G_EEa_, G_EEn_) values as *ν*_ce_ increased. (f) Mean synaptic weights (G_EEa_, G_EEn_, G_IE_, G_EIa_, G_EIn_, and G_II_) for all points maintaining TPA-S as *ν*_ce_ increased. G_EEa_, G_EEn_, and G_IE_ values shown as purple, pink, and red open circles respectively; G_EIa_, G_EIn_, and G_II_ values shown as dark green, light green, and open blue triangles respectively. S.E.M. bars lie beneath the symbols.

#### Loss of excitatory and inhibitory synapses with aging

Peters et al. (Peters et al., 2008) reported a loss of the number of asymmetric and symmetric synapses (excitatory and inhibitory, respectively) in Layers 2/3 of area 46 of rhesus monkey as adult monkeys age: 10% in middle-aged monkeys, and 30% in aged. For the “young monkey model cohort” DRT studies below we represented these respective losses as a 10% decrease in the synaptic weights G_EEa_, G_EEn_, and G_IE_ in middle-aged models, and 30% decrease in these parameters in aged models. For the “young monkey model cohort” DRSTsp studies, we decreased these parameters in a semi-continuous way for the simulated ‘aged’ networks.

### 2.6 Exploring the DRT model parameter space

We conducted two general types of parameter exploration studies for the simulated DRT. The first were sweeps across parameter space using a space-filling Latin Hypercube Sampling (LHS) design, as in (Johnson et al., 1990; Rumbell et al., 2016). The LHS design identified 4200 points (networks) across the parameter space of synaptic weights (G_EEa_, G_EEn_, G_IE_, G_EIa_, G_EIn_, and G_II_), representing semi-random combinations of the synaptic weights homogeneously distributed across the 6-dimensional space (Rumbell et al., 2016) (see Table 2). Bounds of the parameter space were chosen so that the mean firing rate of excitatory neurons in all “young” model networks performing the DRT successfully had mean firing rates of excitatory neurons near 35 Hz, a realistic value for the PFC (see Fig. 5, black curves). We examined the results of network simulations throughout the LHS as the excitatory firing rate parameter *ν*_ce_ varied from 2 to 10 Hz (Fig. 3). All other parameters remained fixed for all simulations. For each of the 4200 parameter combinations in the LHS, the DRT was simulated 7 times to account for variations due to noise in the firing rate equations (Eq. 1), and results were averaged across all repetitions. We identified five main categories of network behavior: maintaining tuned persistent activity (TPA) to any angle orientation until the end of the delay period; maintaining TPA tuned to the original stimulus location at 0° (TPA-S) until the end of the delay period; under-excited networks (either unable to maintain TPA until the end of the delay, or unable to generate a bump of activity at all); over-excited networks (all neurons firing at a similar rate by the beginning of the cue period and for the remainder of the task); and partially over-excited networks (initially maintaining TPA, but becoming over-excited sometime during or after the cue period).

**Table 2.** Parameter ranges for each synaptic weight (in pA·s) used to generate the Latin Hypercube Sample in the DRT model.

| *Synaptic weight* | *Minimum* | *Maximum* |
| --- | --- | --- |
| $G_{EEa}$ | 10 | 80 |
| $G_{EE_n}$ | 30 | 120 |
| $G_{IE}$ | 15 | 60 |
| $G_{EIa}$ | 100 | 240 |
| $G_{EI_n}$ | 100 | 240 |
| $G_{II}$ | 100 | 240 |

**Figure 4:**
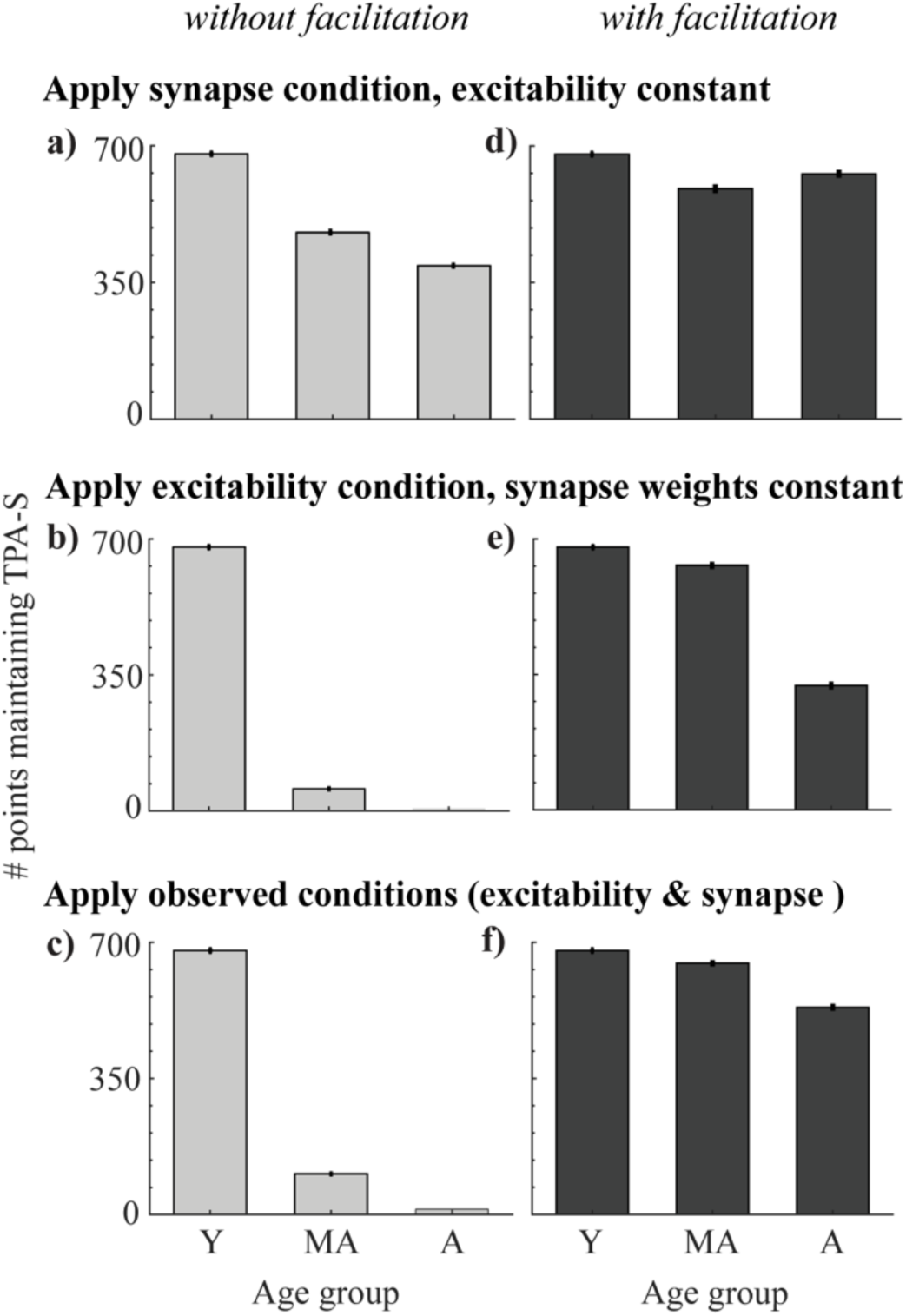
Perturbing the DRT young model cohort. Young cohort defined as networks from the LHS maintaining TPA-S when *ν*_ce_ = 5 Hz (679 networks, dark gray bar in Fig. 3d, and bar for young simulations in all panels in this figure). For middle-aged and aged simulations: (a) Applied the synapse condition to young cohort, holding the excitability parameters constant. (b) Applied the excitability condition, holding synapse parameters constant. (c) Applied both observed data conditions simultaneously (excitability and synapse conditions). (d-f): Analogous perturbations as in (a-c), after adding synaptic facilitation to each network.

**Figure 5:**
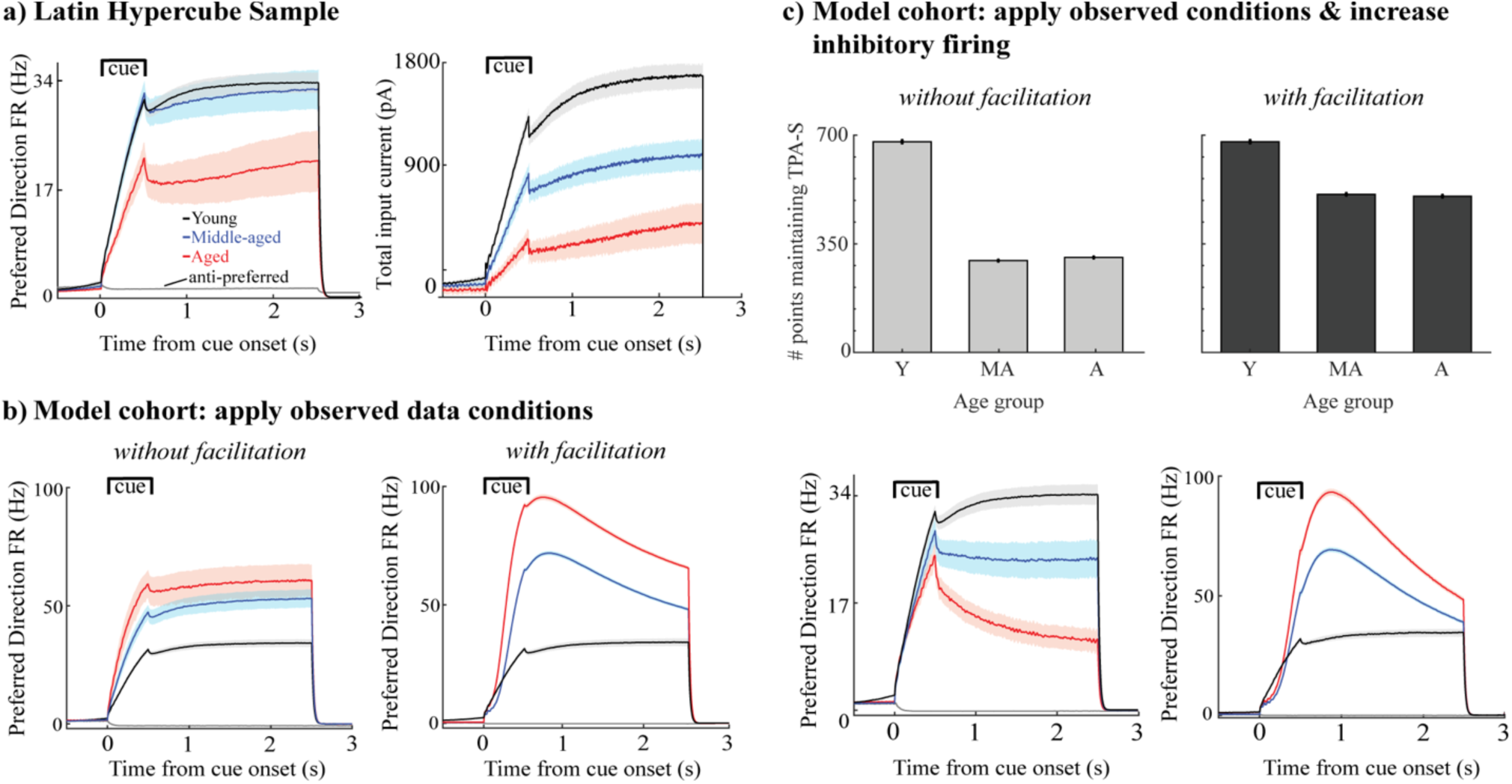
Firing rates and total synaptic current of excitatory neurons in DRT model networks maintaining TPA-S. (a) *Left*, mean FR of excitatory neurons vs. simulated time for all sampled networks maintaining TPA-S when *ν*_ce_ = 5, 7, and 9 Hz (networks shown in Fig. 3e). Shown are neurons with preferred directions matching the 0° stimulus; gray curve shows the mean FR for neurons with a 180° preferred direction, which fired in none of these networks. *Right*, mean total synaptic input current to excitatory neurons with the 0° preferred direction vs. simulation time for the same networks. (b) Mean FR of excitatory neurons for all networks maintaining TPA-S for the model cohort perturbations when the observed data conditions were applied (Fig. 4c and 4g). *Left*: without facilitation. *Right*: with facilitation. (c) *Top row*: DRT model cohort with applied both observed data conditions and increased excitability of inhibitory neurons. *Bottom row*: Mean FR of excitatory neurons for all networks maintaining TPA-S. *Left*: without facilitation. *Right*: with facilitation. In all panels, young, middle-aged, and aged groups are shown in black, blue, and red respectively; shaded regions indicate 95% confidence intervals about each mean.

For the second type of parameter exploration, we first defined a young model cohort as points in the LHS associated with *ν*_ce_ = 5 Hz which maintained TPA-S until the end of the delay period in at least one of the 7 simulation repetitions (shown as the *ν*_ce_ = 5 Hz bar in Fig. 3d). We then perturbed parameters of the young model cohort to simulate the effects of morphological and physiological changes that have been observed with aging, or that we propose as novel possibilities (Fig. 4). Eight kinds of perturbations were made to the young model cohort: (1) The increased excitability condition: increasing *ν*_ce_ from 5 Hz to 7 and 9 Hz, keeping all else constant, to simulate the effect of increasing excitability of individual neurons as monkeys age beyond young adult. (2) The synapse loss condition: reducing the values of G_EEa_, G_EEn_ and G_IE_ by either 10% or 30%, simulating the effect of fewer excitatory and inhibitory synapses onto pyramidal neurons for middle-aged and aged monkeys respectively. (3) Combined increased excitability and synapse loss condition: simultaneously perturbing the excitability and synapse parameters from (1) and (2). Below this perturbation is also called the ‘observed data conditions’. (4) The observed data conditions (combined increased excitability and synapse loss conditions) as in (3), plus increasing the excitability of the inhibitory neurons in the same proportion as in the excitatory neurons: increasing *v*_ci_ from 50 Hz to 70 and 90 Hz for middle-aged and aged simulations respectively. Each of these four perturbations was then repeated after adding short-term synaptic facilitation (as described below) to the networks of the young model cohort. As with the LHS, each young cohort network simulation was repeated 7 times with different randomized seeds and results were averaged.

During *in vivo* recordings, Wang et al. (Wang et al., 2011) found a similar reduction of the firing activity of the DELAY neurons during the cue and delay periods of a DRT with aging, in monkeys performing the task well, and they used computational modeling of the DRT to show that an age-related loss of synaptic strength could account for that reduction. Here we studied if this was still true when the excitability of the excitatory neurons increased at the same time (Fig. 5a). To do this, we examined the mean firing rate of excitatory neurons for the networks in the LHS maintaining TPA-S as *ν*_ce_ increased from young to aged models (for *ν*_ce_ = 5, 7 and 9 Hz). In these simulations we reduced the stimulus scaling factor, I_st_, by 10% and 30% for middle aged and aged networks respectively. This made firing rates during the cue period more consistent with data from Wang et al. (Wang et al., 2011), without significantly affecting firing rates during the delay period (driven primarily by G_EEa_ and G_EEn_).

### 2.7 Simulating memory retention in a DRSTsp – like task

Most of the computational models developed to understand the neural mechanisms underlying working memory are based on delayed memory tasks with just one stimulus to remember during the delay period. A small but growing number of computational studies simulate behavioral tasks that involve the maintenance of several stimuli in memory. Developing models able to maintain the memory of multiple stimuli is challenging, due to interference produced by the simultaneous activation of the many neurons encoding the different stimuli. One way to avoid this problem is to have each stimulus encoded by sparse bumps of activity that does not overlap with firing activity encoding other stimuli (Amit et al., 2003). The levels of inhibition (Edin et al., 2009) and excitation (Wei et al., 2012) in the network are two factors that control the working memory capacity for maintaining multiple stimuli. The balance between them influences the number of stimuli that can be remembered and how bumps fail (fading out vs. merging). Continuous attractor models require fine-tuning of the parameters to control the balance of excitation and inhibition. Such fine tuning can be relaxed by introducing short-term synaptic plasticity. Mongillo et al. (Mongillo et al., 2008) proposed that calcium-mediated synaptic facilitation could be the neural mechanism underlying working memory. Rolls et al. (Rolls et al., 2013) showed that synaptic facilitation can make the system more robust to model parameters. Both found that synaptic facilitation increases the multi-item working memory capacity in a discrete attractor network. Mi et al. (Mi et al., 2017) also found that working memory capacity increased with the time constant of synaptic depression. Some bump attractor models of the DRT have included synaptic facilitation (Itskov et al., 2011; Hansel and Mato, 2013), but in the past it has often been omitted.

We extended the simulation setup of the DRT to create a model of memory retention in a simplified task that is similar to the DRSTsp. Our original bump attractor model had great difficulty maintaining the memory of multiple stimuli simultaneously, due to interference between the stimulus responses (Amit et al., 2003; Barak and Tsodyks, 2014) and because during the first cue and delay periods, all neurons not involved in the bump activity were highly inhibited. Thus, when introducing subsequent stimuli, the stimulus input current (equal for all stimuli) was insufficient to induce subsequent bumps of firing activity. This difficulty was addressed by adding short-term synaptic facilitation in the excitatory-to-excitatory synaptic connections. As in (Itskov et al., 2011), we assumed that the overall behavior of the synapses was facilitating, and modified the first two equations in Eq. 3 as

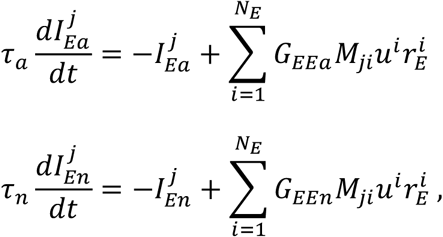

with the facilitation variable *u* given by

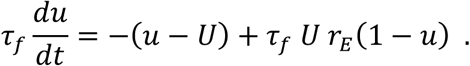

Here *u* is a utilization variable defining the fraction of total neuronal resources that each spike is able to access, representing increased calcium levels at presynaptic terminals (Mongillo et al., 2008). Without firing, *u* decays to its baseline value *U* with time constant *τ*_*f*_. In all our simulations *U* = 0.001and *τ*_*f*_ = 1500 ms.

Behavioral data showed a maximum DRSTsp score of 3.61 for one of the middle-aged subjects (Fig. 1b). The cognitive score recorded is the recognition span: the mean number of stimuli the monkey identified correctly before making a mistake. Thus, we designed our computational task so that perfect performance gave a simulated DRSTsp score (hereafter called *DRST*_*sim*_) of four, corresponding to successful encoding of four stimuli equally spaced around the ring from the DRT (Fig. 6a). Our DRSTsp model simulated a ‘pre-cue’ period of 1 s, followed by alternating cue and delay periods (lasting 0.5 s and 2 s respectively). The pre-cue and delay periods represented times in the real DRSTsp when the screen was down while appropriate wells were baited. In each subsequent cue period, a new stimulus was presented in addition to any stimuli that were presented before. In the first cue period a stimulus was presented at 0°; in the second, stimuli were presented at 0° and 90°. In the third cue period stimuli were presented at 0°, 90° and −90°, and in the fourth stimuli were presented at 0°, 90°, −90° and 180°. Stimuli were identical each time at each presentation, with I_st_ = 40000 pA and *c* = 20. After the first cue period, if a network simulation encoded all stimuli during cue period *n* and maintained all *n* – 1 stimuli throughout the previous delay period (by maintaining a bump for each stimulus with maximum FR of at least 5 Hz), we assumed that the novel stimulus was identified correctly. In this way, our task models the ability of our network to retain the presented stimuli, but does not model the process of choosing which stimulus among all presented is the novel one. Our task also assumes that each trial period lasts the same amount of time, whereas monkeys performing the DRSTsp terminate each successful trial by moving the disk corresponding to their choice before the maximum trial time expires. Sample networks with *DRST*_*sim*_ = 4, 3, 2, and 1 are shown in Figure 6b.

**Figure 6:**
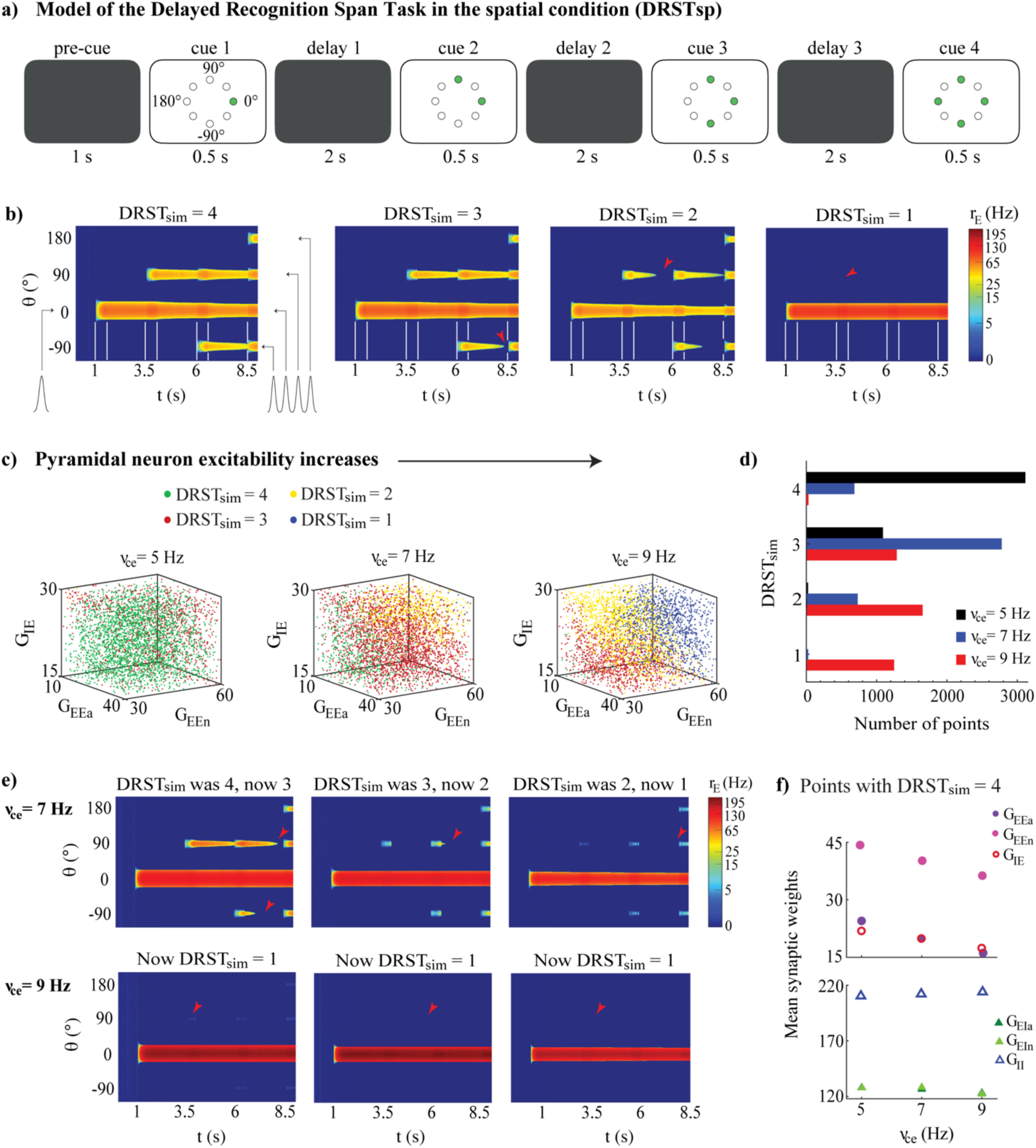
Memory retention model of the DRSTsp shows impairment under neuronal hyperexcitability. The ability to maintain the memory of a successively increasing number of stimuli strongly depended on the excitability of individual pyramidal neurons, but changes in the synaptic weighs partially compensated for the increased neuronal excitability. (a) Simulated task included a pre-cue period, with four cue periods separated by three delay periods. Green circles indicate the stimuli presented in each cue period. (b) Examples of DRSTsp model output, with FR shown in color (log scale) for neurons labeled by their preferred direction vs. simulated time. From left to right: *DRST*_*sim*_ = 4, optimal performance, all four stimuli were encoded correctly and recalled; *DRST*_*sim*_ = 3, where the first three stimuli were encoded correctly, but persistent activity representing the third was lost during the subsequent delay period; and *DRST*_*sim*_ = 2 and *DRST*_*sim*_ = 1 where only the first two or one stimuli were encoded correctly and recalled. Cue periods shown between white lines; arrowheads indicate when persistent activity was lost. (c) DRSTsp model output for 4200 points of the parameter space LHS, shown in 3D projections across the (G_EEa_, G_EEn_, G_IE_) subspace as *ν*_ce_ increased. Points with *DRST*_*sim*_ = 4, 3, 2, and 1 are shown as green, red, yellow, and blue dots respectively. (d) Histogram of *DRST*_*sim*_ values in the LHS as *ν*_ce_ increased (*ν*_ce_ = 5, 7, and 9 Hz shown as black, blue, and red bars respectively). (e) The first three simulations from panel (b), repeated for *ν*_ce_ = 7 (top row) and 9 Hz (bottom row). (f) Mean synaptic weights for all LHS points with *DRST*_*sim*_ = 4 for *ν*_ce_ = 5, 7, and 9 Hz. Top: G_EEa_, G_EEn_, and G_IE_ shown as purple, pink, and red open circles. Bottom: G_EIa_, G_EIn_, and G_II_ shown as dark green, light green, and open blue triangles. S.E.M. bars lie beneath the symbols.

With this new *DRST*_*sim*_ established, we again used LHS to explore how task performance varied across the parameter space of the six synaptic weights (G_EEa_, G_EEn_, G_IE_, G_EIa_, G_EIn_, and G_II_; see Table 3) and for *ν*_ce_ = 5, 7 and 9 Hz. As for the DRT, each LHS had 4200 points (Fig. 6c); we chose the bounds of the parameter space so that all 4200 networks encoded at least the first stimulus for all three *ν*_ce_ values. Simulations were performed only once, since synaptic facilitation made the network performance much less sensitive to random fluctuations.

**Table 3.**
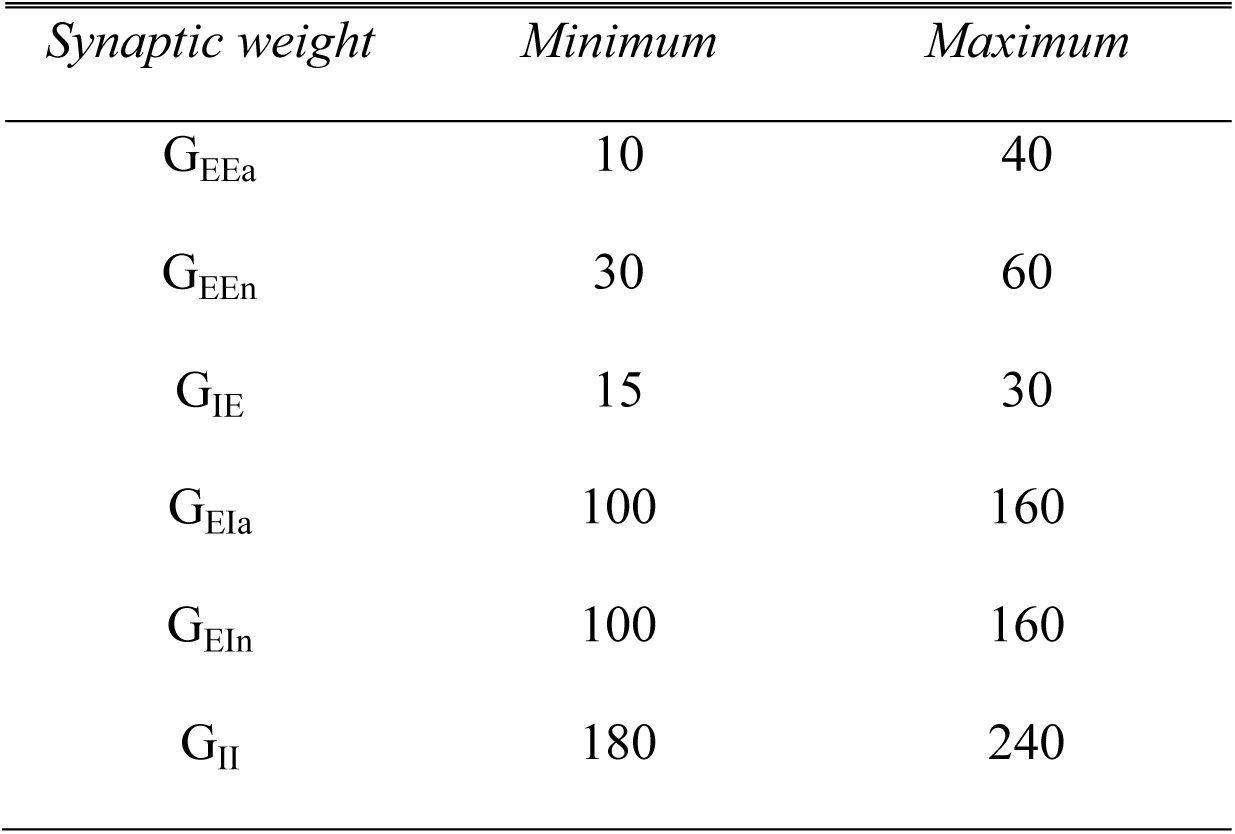
Parameter ranges for each synaptic weight (in pA·s) used to generate the Latin Hypercube Sample in the DRSTsp model.

We also defined a young monkey model cohort for the DRSTsp model to be a group of 40 randomly chosen points in the LHS with *ν*_ce_ = 5 Hz (Fig. 6d, young case: black bars), half with *DRST*_*sim*_ = 4 and the other half with *DRST*_*sim*_ = 3. We then “aged” the networks in this cohort by applying increased excitability and synaptic loss conditions, and increasing the excitability of the inhibitory neurons, as follows. We set bounds for *ν*_ce_ just beyond our young and aged values (4.5 and 9.5 Hz) and generated 40 randomly distributed values between them. We then defined two piecewise linear functions that interpolated the synapse loss conditions from the DRT (0%, 10% and 30% loss when *ν*_ce_ = 5, 7, 9 Hz respectively, slope 5% loss/Hz when *ν*_ce_ < 7 Hz and 10% loss/Hz when *ν*_ce_ ≥ 7), and computed a corresponding level of synapse loss for each of the 40 *ν*_ce_ values. We computed corresponding values for *v*_ci_ by multiplying each *ν*_ce_ value by 10, consistent with the relationship between these two parameters elsewhere in our study. We computed an associated age for each *ν*_ce_ value similarly, interpolating the mean age for the three empirical groups (8.9, 18.2, and 24.7 years for young, middle aged, and aged groups) when *ν*_ce_ = 5, 7, 9 Hz respectively. We induced the increased excitability and synapse loss conditions of these 40 values separately and together, plus increased excitability of inhibitory neurons, and then computed the corresponding *DRST*_*sim*_ for each transformed member of the young cohort (6400 simulations in all: 160 for each point in the cohort). We then calculated the mean *DRST*_*sim*_ for each simulated age group (young group, 7.5 < years < 13; middle-aged group, 13 ≤ years < 21; and aged group, 21 < years < 26). To summarize the change in performance across the simulated age span (Fig. 7), we calculated the relative change in the mean *DRST*_*sim*_ from the mean ages of the young simulations to the aged simulations (9.9 and 24.2 years respectively) under each perturbation condition.

**Figure 7:**
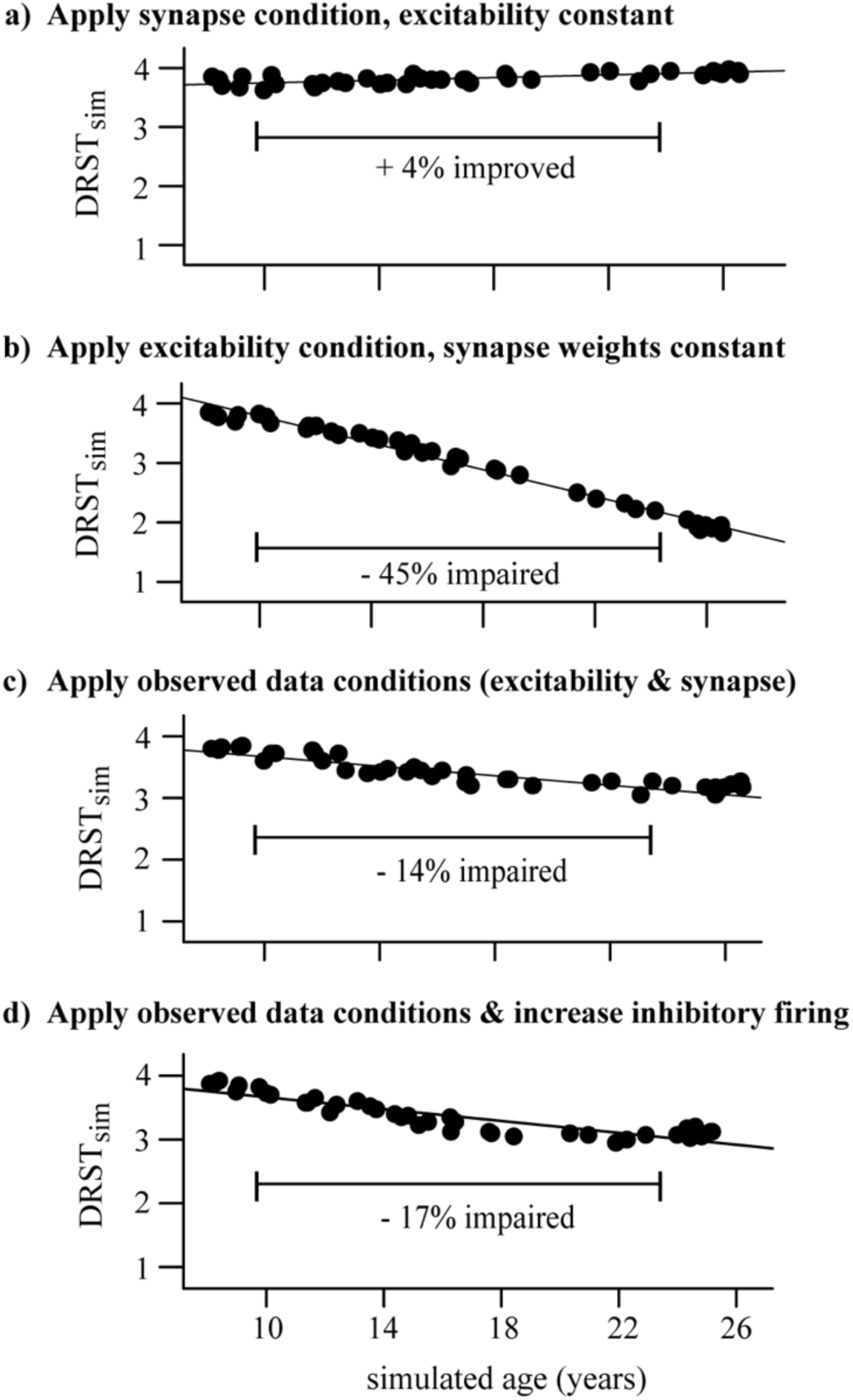
Perturbing the DRSTsp young model cohort. Each dot indicates the mean *DRST*_*sim*_ of the 40 young cohort points. (a) Synapse condition only; slight improvement (+4%) from the young to the aged simulations. (b) Excitability condition only; strong impairment (−45%) from young to aged. (c) Applied both observed conditions simultaneously, resulting in just a −14% impairment. (d) Applied both observed data conditions and increased excitability of inhibitory neurons; −17% impairment.

### 2.8 Statistics

All analyses were performed in RStudio 1.1.463 (RStudio Team, 2015; Mangiafico, 2016). We treated age as a continuous variable in most analyses, using standard linear regression as well as generalized linear mixed-effects models (GLMMs) to explore relationships between output variables vs. dependent variables. Because electrophysiological data were collected from several neurons per subject, the subject was treated as a random effect blocking factor in relevant analyses (Grafen and Hails, 2002; Darian-Smith et al., 2013). For the GLMMs, marginal *R*^2^ captured the variance explained by fixed factors; conditional *R*^2^ captured the variance explained by the whole model including the random effects (Nakagawa and Schielzeth, 2013). In analyses with DRSTsp score (one data value per subject) versus physiological variables, physiological variables were first averaged within each subject.

Age was treated as a categorical variable (young, middle-aged, and aged) when analyzing relationships firing rate vs. age (Fig. 1e). Specifically, we used two-way repeated measures ANOVA (including subject as a random effect) to determine whether there were differences in FR vs. age group, with injected current as the repeated measure. Before performing the ANOVA, quantile-quantile plots were used to identify extreme outliers from normality for each age group and level of injected current, and then Shapiro-Wilk’s test of normality was applied. Post-hoc analyses were conducted using Tukey-adjusted comparison of estimated marginal means. For all analyses, the significance level was α = 0.05.

## 3. RESULTS

### 3.1 Rhesus monkeys show DRSTsp impairment and increased AP firing rates of dlPFC pyramidal neurons with aging

We examined relationships between performance on the DRSTsp, age of the monkeys, and physiological variables assessed (Fig. 1). Among the 39 subjects analyzed for this study, 36 of them completed behavioral testing on the DRSTsp (6 young, 18 middle-aged, and 12 aged). There was a significant decrease in performance on the DRSTsp with age (linear regression, DRST = −0.032*Age + 2.97, *R*^2^ = 0.18, F_(1,34)_ = 7.43, p = 0.0101; Fig. 1b) replicating past findings on impaired working memory with age in monkeys (Herndon et al., 1997; Moore et al., 2006; Wang et al., 2011; Moore et al., 2017). *In vitro* slices were prepared from the dlPFC of the same monkeys that were tested behaviorally for whole cell patch clamp recordings. Data from 324 neurons met our criteria for inclusion. Our statistical analysis used random effects modeling to control for the variation of physiological variables within monkeys (Materials & Methods). Input resistance (R_n_) increased with aging (R_n_ = 4.02*Age + 89.1 M*Ω*, marginal *R*^2^ = 0.08, conditional *R*^2^ = 0.20, p < 0.0001; Fig. 1c), while resting membrane potential did not (p = 0.11). There was a significant positive relationship between input resistance and AP firing rate at all levels of current injection (p < 0.00016 and conditional *R*^2^ ≥ 0.21 for each level, Fig. 1d).

Next, we compared depolarizing current step-evoked AP firing rate in neurons from young, middle-aged, and aged subjects (Fig. 1e). Before using the two-way repeated measures ANOVA, outliers from normality were removed for AP firing rate vs. current injection level for each age group. There was a significant effect of age group on firing rate (F_(2,22)_ = 8.63, p = 0.0017) but no significant interaction between age group and injected current (F_(2,799)_ = 2.76, p = 0.064). Firing rates were significantly higher for the aged vs. young monkeys (Tukey-adjusted comparison of estimated marginal means, t_(22)_ = 4.14, p = 0.0012); firing rates for middle-aged monkeys did not differ from the other two groups (young vs. middle-aged, t_(22)_ = 2.23, p = 0.072; middle-aged vs. aged, t_(22)_ = 1.99, p = 0.139; Fig. 1e). Hereafter we refer to this increased firing rate of current step-evoked APs as hyperexcitability of pyramidal neurons.

Given significant relationships between aging and both DRSTsp performance and the physiological variables, we next examined whether both aging and physiological variables might together affect DRSTsp performance. Since R_n_ and firing rate were significantly related at each injection level, we only included a single physiological variable in the analysis: firing rate in response to the 130 pA current injection. There was a significant negative linear relationship between DRSTsp span and firing rate (DRSTsp = −0.055*FR + 2.95 Hz, *R*^2^ = 0.19, F_(1,33)_ = 7.71, p = 0.0090, Fig. 1f). To examine the effect of age and firing rate on DRSTsp span, we fitted a regression model that included age, firing rate, and their interaction (*R*^2^ = 0.26, F(3,31) = 3.68, p = 0.0225). The negative effect of the firing rate on DRSTsp was significant (p = 0.0441) and it did not depend on the age of the animal (age by firing rate interaction effect p = 0.1340). The effect of age alone was not significant either (p = 0.0904). These analyses lend support to the hypothesis that firing rate affects DRSTsp performance more than age does, though this support is somewhat tenuous for the following reasons. First, the delay between DRSTsp assessment and physiological recording of neurons from these valuable subjects can last several months to a year or more, and may mask a relationship between these variables. Second, some of these subjects were employed in longitudinal studies for other projects and tested on the DRSTsp a few times; we cannot rule out a practice effect in these subjects. Thus, we turned next to computational modeling to create a controlled environment for exploring how changes in AP firing rates (independent of a subject’s age) might affect performance on spatial working memory tasks.

### 3.2 Model 1: The Delayed Response Task (DRT)

#### 3.2.1 An optimal level of pyramidal neuronal excitability maximized successful performance in the DRT model

Here we used computational modeling to explore how these empirical results of physiological changes and a past study on reduced synapse counts with aging might together impact cognitive function in middle-aged and aged monkeys. We chose a continuous bump attractor network model which has been used by several previous research groups (Compte et al., 2000; Wang et al., 2011; Wang et al., 2013; Wimmer et al., 2014). The network model simulates an empirically measured behavior (the oculomotor Delayed Response Task administered to macaque monkeys) through a simple but elegant model network that can be manipulated in specific ways (Fig. 2; Materials & Methods). During the cue period of the DRT simulations, all excitatory neurons received an external input current, tuned to 0° direction, proportional to their orientation preference (Fig. 2d). In the subsequent delay period, the main input to model neurons came through recurrent activity within the network. If persistent neural activity is maintained at the end of the delay period—in the form of a bump-shaped FR profile for a subset of the excitatory neurons—then the orientation angle corresponding to the center of the bump of activity at the end of the delay is assumed to represent the stimulus location encoded by the network (Fig. 2e).

Varying model parameters created a distinct network configuration, and each simulation represented a model DRT trial. Successful DRT trials maintained TPA (and in particular TPA-S) until the end of the delay period (Fig. 2e and 3a_2_). Failed DRT trials can occur in several ways, including under-excitation, a loss of TPA during the cue or delay periods (Fig. 3a_1_); generating TPA early in the cue or delay periods but ending with full network over-excitation (Fig. 3a_3_); and an over-excited network throughout the simulation (Fig. 3a_4_).

An important feature of the bump attractor model we chose to use (Wimmer et al., 2014) is the ability to control the *f-*I curve of individual neurons by two simple parameters. Adjusting the parameter *ν*_ce_ was sufficient to fit the *f*-I curves of excitatory model neurons to the mean *in vitro* AP firing rates of pyramidal neurons reported above (Fig. 3b and Eq. 2). Young, middle aged, and aged *f*-I curves used *ν*_ce_ values of 5, 7, and 9 Hz respectively. These fits were accurate for low input current, but became more excitable than the empirical *f-I* curves for high inputs. However, our results did not depend on the specific shape of the *f-I* curves: similar results were found when using less excitable analogues of the firing rate function (Supplementary Fig. 1). Synaptic weight parameters represented the number of excitatory and inhibitory synapses onto excitatory (G_EEa_, G_EEn_, and G_IE_) and inhibitory (G_EIa_, G_EIn_, and G_II_) neurons. Varying these synaptic weights allowed us to model published data showing an age-related reduction in the excitatory and inhibitory synapses with aging in the neuropil of layer 2/3 of rhesus monkey dlPFC (Peters et al., 2008). In a series of simulations, we kept *ν*_ce_ fixed at distinct values and then used the Latin Hypercube Sampling (Materials & Methods) to choose points throughout the parameter space generated by the six synaptic weights. Each point was simulated seven times, as if administering multiple DRT trials to each virtual monkey. We examined whether points across the parameter space maintained TPA throughout the delay period, or had a different outcome (Fig. 3c).

Increasing the excitability of model excitatory neurons strongly affected the ability of the DRT model networks to maintain TPA. The 3D plots in Figure 3c summarize DRT model results for 4200 points across the parameter space of synaptic weights as *ν*_ce_ increased. Blue circles represent ‘under-excited’ networks, where no TPA was detected at the end of the delay period (e.g., Fig. 3a_1_). Green circles represent networks that maintained TPA for the entire delay period (e.g., Fig. 3a_2_)—corresponding to ‘virtual monkeys’ that encoded some stimulus location. Remaining circles represent partially over-excited networks (yellow; e.g., Fig. 3a_3_), which started out with TPA that led to firing in all neurons, or were already over-excited networks during the fixation period before the cue ever appeared (red; e.g., Fig. 3a_4_). Below we focus on the green points, which maintained TPA throughout the delay period.

The greatest likelihood of maintaining TPA, and in particular TPA-S, across the synaptic weight parameter space occurred for intermediate values of *ν*_ce_ – neither too low nor too high (Fig. 3d). The optimal value among those we tested was *ν*_ce_ = 5 Hz. Values of *ν*_ce_ below this optimum led to many networks across parameter space that were under-excited; above the optimum many networks were over-excited (Fig. 3c). In either case, the networks were much less likely to encode the stimulus successfully until the end of the delay period, and this reduced likelihood was visible even in our ‘middle-aged’ networks (*ν*_ce_ = 7 Hz). Thus, if the excitability of single pyramidal neurons increased above the optimal value without complementary changes in other parameters, the network’s ability to maintain tuned persistent activity, in particular to the stimulus location— and successful working memory performance—was impaired. These findings also held with the less excitable form of the *f*-I curves fit to the data (supplemental Figs. 1b and 1d).

#### 3.2.2 Synaptic changes partially compensated for increased pyramidal neuron excitability in successful DRT model networks

Points across parameter space that did maintain TPA-S as *ν*_ce_ increased had lower values of the excitatory synapse weights for excitatory neurons (G_EEa_ and G_EEn_) and higher values of the inhibitory synapse weights (G_IE_) for these neurons (see Fig. 3f; standard error bars lie beneath the symbols). Figure 3e shows the mean values of each synaptic weight for all sampled points maintaining TPA-S as *ν*_ce_ increased. There was no clear relationship between *ν*_ce_ and the synaptic weight parameters onto interneurons (G_EIa_, G_EIn_, and G_II_). However, excitatory parameters G_EEa_ and G_EEn_ decreased noticeably as *ν*_ce_ increased before leveling off after the middle-aged *ν*_ce_ value, while G_IE_ increased in a nearly linear fashion. To maintain TPA in the DRT model (particularly when tuned to the stimulus location), the increased excitability of excitatory neurons was compensated by altering synaptic weights onto those same neurons: with less synaptic excitation and more synaptic inhibition. Yet despite the compensatory synaptic weight changes, fewer networks were able to maintain TPA-S as *ν*_ce_ increased (Fig. 3d). See also supplemental Figure 1c for the generalized *f*-I curve.

These LHS simulations revealed that network function can be maintained somewhat as parameters vary freely to compensate pyramidal neuron hyperexcitability with aging, but the predicted compensation is not consistent with the empirically observed *loss* of both excitatory and inhibitory synapses on pyramidal neurons with aging. To explore how concomitant changes to firing rates and synapses might affect network function, we first considered all networks maintaining TPA-S when *ν*_ce_ = 5 Hz as corresponding to a simulated cohort of cognitively healthy young monkeys (the 679 model networks with *ν*_ce_ = 5 Hz in Fig. 3d, called the ‘young model cohort’). We then examined the performance of the model networks in this cohort on the DRT after perturbing parameters in proportions that matched empirically observed changes with aging (Fig. 4). First is the hyperexcitability for pyramidal neurons *in vitro* described above and in our past work (Chang et al., 2005; Coskren et al., 2015). Second, Peters et al. (Peters et al., 2008) reported a ∼10% loss of excitatory and inhibitory synapses in the rhesus dlPFC neuropil by middle-age, and a ∼30% loss of both synapse types in aged subjects. This finding is consistent with other studies of synapse and spine loss with aging in the rhesus dlPFC (Hof et al., 2002; Duan et al., 2003; Young et al., 2014).

Accordingly, we induced “middle-aged” and “aged” conditions for pyramidal neuron excitability in the young cohort by increasing *ν*_ce_ to 7 and 9 Hz respectively. To model synapse loss, we reduced G_EEa_, G_EEn_ and G_IE_ in the young model cohort by 10% and 30% respectively to create either a middle-aged or aged synapse condition. We then induced the two conditions separately and together (forming the ‘observed data conditions’), and determined whether each transformed model still maintained TPA-S during DRT simulations (Fig. 4a-c). A similar synapse loss condition for excitatory synapses was modeled by Wang et al. (Wang et al., 2011) to explain an age-related decrease in firing rates during *in vivo* recordings of the DELAY neurons for monkeys successfully performing the DRT. Our study builds on Wang et al. (Wang et al., 2011) by adding inhibitory synapse loss and *f*-I curve changes with aging, and examining how these alterations affect task performance for a range of parameter values.

DRT performance of the young model cohort was much more sensitive to the excitability aging condition than to the synapse condition (Fig. 4a-b). Even under the strongest (aged) synapse condition, over half the networks still maintained TPA-S (Fig. 4a). In contrast, under the excitability condition most of the networks had already lost their capacity for TPA-S by middle age (Fig. 4b). In the parameter space exploration above, pyramidal neuron hyperexcitability was partially compensated by decreasing excitatory synapse weights and increasing inhibitory ones. Thus, while Peters et al. (Peters et al., 2008) found that inhibitory synapse counts actually *decrease* with aging, we might still expect a partial recovery of function when the young cohort underwent the excitability and synapse aging conditions simultaneously (‘observed data conditions’, Fig. 4c). However, this was not the case, as almost none of the middle aged and aged models maintained TPA-S after these perturbations. This does not seem to match reality, since Wang et al. (Wang et al., 2011) noted no difficulty in identifying middle-aged and aged monkeys who perform the DRT successfully, and impairment on the more difficult DRSTsp is noticeable but not severe by middle age (Fig. 1b).

Recent studies have shown that including short-term synaptic facilitation in discrete and continuous attractor models can reduce the fine-tuning needed to maintain performance (Mongillo et al., 2008; Itskov et al., 2011; Rolls et al., 2013). We added short-term synaptic facilitation to our middle-aged and aged DRT model networks (same parameter values in both groups), as a baseline exploration of how this mechanism might affect DRT performance with aging. The results were qualitatively similar to those without facilitation, except the model cohort networks were affected less by the aging perturbations (Fig. 4d-f). Under the synapse condition, almost all networks maintained TPA-S both for middle-aged and aged simulations, with even a slight increase from middle-aged to aged. Networks were still more sensitive to the excitability aging condition than to the synapse condition (Fig. 4d-e), but under the excitability condition nearly all the middle-aged and about half the aged networks maintained TPA-S. Applying both observed aging conditions together with facilitation led to more networks maintaining TPA-S (Fig. 4f), suggesting that synaptic facilitation can boost DRT performance.

#### 3.2.3 Examining firing rates during the simulated DRT

Wang et al. (Wang et al., 2011) examined *in vivo* firing rates of area 46 (dlPFC) pyramidal neurons in rhesus monkeys across the adult age span while they performed the DRT. For monkeys performing the task well, they found the firing rate of DELAY neurons during the cue and delay periods decreased with aging, both for neurons whose preferred and anti-preferred directions matched the cue presented. In our sampling of parameter space (Fig. 3) we found that DRT model networks maintaining TPA-S—representing monkeys that perform the DRT successfully—had lower firing rates during both the cue and delay periods as *ν*_ce_ increased (Fig. 5a, left). This was particularly true for neurons whose preferred direction matched the stimulus, with a substantial decrease from young and middle-aged to aged model networks. This seemed counterintuitive at first, since increasing *ν*_ce_ makes individual excitatory neurons more excitable in response to a given input current (Fig. 3b). Indeed, hyperexcitability of individual neurons often did lead to over-excitation throughout the network (yellow and red points in Fig. 3c). However, the networks that maintained TPA-S as *ν*_ce_ increased compensated for the increased excitability with a reduction in G_EEa_ and G_EEn_ and increase in G_IE_ (Fig. 3f), leading to lower firing rates. Cue period FR in our model was driven much more by the stimulus current than by synaptic weights, and decreased less across the age groups than the delay period FR did. As such, reducing the stimulus current in the sampled middle aged and aged networks was necessary to match the Wang et al. (Wang et al., 2011) cue period results (Materials & Methods). This change is reasonable, since external inputs *in vivo* are mediated by synapses whose numbers likely also decrease with aging as in Peters et al. (Peters et al., 2008). As a whole, these parameter space explorations show a way to unify our *in vitro* data and *in vivo* experiments, provided that the total input current to pyramidal neurons during the DRT, I_E_, is lower in middle-aged and aged monkeys than in young monkeys (Fig. 5a, right).

We also examined firing rates in the model cohort simulations perturbed by the observed data conditions with and without facilitation (Fig. 4c and 4f). The main distinction between the parameter space sampling simulations and the cohort simulations was how synaptic inhibition onto the excitatory neurons, G_IE_, varied as the excitability parameter (*ν*_ce_) increased: G_IE_ increased in the parameter space explorations but decreased in the cohort simulations due to observed inhibitory synapse loss with aging. Indeed, the weakened inhibitory connections in the middle-aged and aged cohorts led to a FR *increase* during the cue and delay periods of those networks compared to the young cohort (Fig. 5b, left), opposing the Wang et al. (Wang et al., 2011) results. The increase was even greater when synaptic facilitation was included (Fig 5b, right).

Our young cohort simulations did not match data from the *in vitro* physiology, synapse counts, and *in vivo* physiology findings simultaneously, suggesting that one or more age-related changes to the system may be missing from our current model. We simulate one such possibility here, to demonstrate how the current model can help us test hypotheses and incorporate new data as they become available. Specifically, our parameter space explorations showed that an overall increase in inhibition onto the excitatory neurons can compensate for increased firing rates of excitatory neurons. Thus, we examined how network function would change if the observed data conditions were accompanied by a comparable age-related increase of inhibitory neuron firing rates—an empirical question whose answer is not yet known. In models without synaptic facilitation, more middle-aged and aged networks maintained TPA-S when inhibitory neuron firing rates increased with aging (Fig. 4c vs. Fig. 5c, top left). Firing rates of excitatory neurons in those simulations also decreased with aging as observed *in vivo*, consistent with the Wang et al. data (Wang et al., 2011; Fig. 5c, bottom left). However, results differed for models that included synaptic facilitation: fewer middle-aged and aged networks maintained TPA-S when inhibitory neuron firing increased (Fig. 4f vs. Fig. 5c, top right), while (as before) firing rates of excitatory neurons during the DRT increased with aging (Fig. 5c, bottom right). These results suggest that our DRT model might need further refinement to reflect reality better – for example, by also incorporating synaptic depression (Mi et al., 2017) or changes in synaptic plasticity with aging. Regardless, our model allows us to quantify how much the known age-related changes at the single cell level affect cognitive performance, and to predict the extent to which other relevant processes might contribute to working memory impairment with aging.

### 3.3 Model 2: Working memory retention in a DRSTsp-like task

#### 3.3.1 Pyramidal neuron hyperexcitability leads to reduced memory capacity in simulated DRSTsp networks

These DRT modeling results add to the wide range of literature using the bump attractor model to study spatial working memory, but the DRT was not administered to the monkeys from this study. Spatial working memory capacity in these monkeys was assessed with the more complex DRSTsp. To see what predictions our model might have for these data, we extended our DRT model to simulate memory retention in a task with three important similarities to the DRSTsp: (1) the network encoded several stimuli simultaneously, with stimuli that are (2) spatially distinct; and (3) presented to the network successively over time. The changes made to extend the DRT model simulations to a task like the DRSTsp are shown in Figure 6a, and a simulated cognitive score (*DRST*_*sim*_) was assigned to the outcome of each simulation. Examples of model networks with a *DRST*_*sim*_ of 4 (perfect performance) down to 1 are shown in Figure 6b. The second network achieved a *DRST*_*sim*_ of 3 because it encoded stimuli 1, 2, and 3 correctly but lost persistent activity corresponding to stimulus 3 in the subsequent delay period (red arrowhead). Likewise, the third network (*DRST*_*sim*_ = 2) lost persistent activity corresponding to stimulus 2 before stimulus 3 was applied; the final network (*DRST*_*sim*_ = 1) encoded stimulus 1 properly but not stimulus 2.

We examined how *DRST*_*sim*_ varied across the parameter space of synaptic weights for the young, middle-aged and aged network models (*ν*_ce_ = 5, 7, and 9 Hz; Fig. 6c). Networks achieving a *DRST*_*sim*_ of 1 (worst), 2, 3, and 4 (best) are shown respectively as blue, yellow, red, and green. As in the DRT, the value of *ν*_ce_ strongly affected the ability of the DRSTsp model networks to maintain persistent activity tuned to the location of the different stimuli as the number of delay periods and unique stimuli increased. There were many fewer green points in the parameter regime explored, and more blue points, as *ν*_ce_ increased. (As *ν*_ce_ increased, networks with the best *DRST*_*sim*_ performance had lower values of G_EEa_, G_EEn_ and G_IE_ than those shown here.) Figure 6d summarizes these counts within our regime, showing a clear leftward shift in the distribution of *DRST*_*sim*_ across the parameter space (poorer performance) as *ν*_ce_ increased.

Figure 6e shows how the model networks of Figure 6b were affected by increasing *ν*_ce_ from 5 (in Fig. 6b) to 7 and 9 Hz. In general, as *ν*_ce_ increased, the initial 0° stimulus was encoded by a stronger bump (higher firing rate) for the entire DRSTsp simulation, and subsequent stimuli encoded by comparatively weaker bumps. This led to an under-excitation of the network during the second and third delay periods, losing the ability to maintain later stimuli. An example can be seen in the second column of Figures 6b and 6e. For the young monkey case (Fig. 6b, *ν*_ce_ = 5 Hz), the maximum firing rate of the first bump was around 75 Hz; this and the width of the first bump decreased slightly as the simulation continued. Also, as noted above, persistent activity for stimulus 3 was lost during the third delay period. For the middle-aged case (Fig. 6e top row, *ν*_ce_ = 7 Hz), the maximum firing rate of the first bump maintained a maximum firing rate near 150 Hz and a similar width throughout the simulation, but the second bump was lost early in the subsequent delay period. This trend was even stronger for the aged case (Fig. 6e bottom row, *ν*_ce_ = 9 Hz), where the first bump remained wide with a maximum firing rate of 250 Hz throughout the simulation but none of the subsequent stimuli were encoded.

To see how synaptic parameters compensated to maintain DRSTsp performance as *ν*_ce_ increased, we looked at the mean values of the synaptic weights for all networks with perfect task performance (*DRST*_*sim*_ = 4). values of both the excitatory and inhibitory synaptic weights for excitatory neurons were lower as *ν*_ce_ increased (Fig. 6f top; standard error bars lie beneath the symbols). Excitatory synaptic weights for inhibitory neurons decreased and inhibitory synaptic weights increased as *ν*_ce_ increased. (Fig. 6f bottom; standard error bars lie beneath the symbols). Overall, for the DRSTsp model, both panels showed that synaptic excitation as well as inhibition to excitatory neurons decreased to partially compensate for the increased excitability of individual excitatory neurons.

#### 3.3.2 “Aging” the DRSTsp model networks leads to working memory impairment as observed empirically

As with the DRT model, we defined a simulated “young monkey cohort” for the DRSTsp model and “aged” the model networks in this cohort by applying increased excitability and synaptic loss conditions (Fig. 7), choosing 40 levels of parameter perturbations throughout the simulated age span. We induced the two observed data conditions separately and together—and to match Figure 5c top right, also increased the excitability of inhibitory neurons. We then computed the corresponding *DRST*_*sim*_ for each transformed member of the young cohort. To summarize the change in performance across the simulated age span, we calculated the relative change in the mean *DRST*_*sim*_ from the young to the aged models under each perturbation condition. This facilitated comparison between performance on the simulated task (four stimuli spaced evenly around a ring), and the real DRSTsp (18 possible stimuli, organized spatially into three rows of six wells each).

All four cases showed a strong linear relationship between *DRST*_*sim*_ and parameter perturbations. As in the DRT model above, the DRSTsp performance of the young model cohort was much more sensitive to the excitability condition (Fig. 7b) than to the synapse condition (Fig. 7a). While the synapse condition showed a 4% *improvement* with aging for the DRSTsp model (*R*^2^ = 0.98, p < 0.0001), the excitability-only condition showed a 45% decrease in *DRST*_*sim*_ with aging (*R*^2^ = 0.98, p < 0.0001). Applying the two observed conditions together led to a 14% impairment with aging (Fig. 7c; *R*^2^ = 0.82, p < 0.0001); applying the observed data conditions plus increasing the excitability of inhibitory neurons led to a 17% impairment (Fig. 7d; *R*^2^ = 0.81, p < 0.0001). Thus, both cases of applying the observed data conditions were comparable to the 20% impairment exhibited by the empirical data (Fig. 1b). These results are also consistent with Figure 6f: the empirically-observed decrease in synaptic excitation and inhibition to excitatory neurons partially compensated the increased excitability of the excitatory neurons, so that *DRST*_*sim*_ impairment with aging was less severe. The model also suggests that hyperexcitability of inhibitory neurons would impact performance on the DRT more than it would the DRSTsp.

## 4. DISCUSSION

This study used computational modeling to explore how single pyramidal neuron hyperexcitability and excitatory and inhibitory synapse loss observed with aging in layer 3 of the rhesus monkey dlPFC might contribute to spatial working memory impairment. First, we presented new empirical data from rhesus monkeys across the adult age span, consistent with past studies, showing that working memory impairment and increased FRs of neurons occur in some middle-aged monkeys. This replicated previous studies showing a decline in cognitive performance from young to aged monkeys (Herndon et al., 1997; Moore et al., 2003; Moore et al., 2006; Konar et al., 2016; Motley et al., 2018), and a concomitant increase in R_n_ and FR of dlPFC pyramidal neurons (Chang et al., 2005; Coskren et al., 2015). These data motivated the modeling, and were used to constrain a bump attractor network model of the DRT (Wimmer et al., 2014) and our network model of working memory retention in the DRSTsp. These models predict that the observed age-related firing rate increase and synapse loss in rhesus dlPFC (Peters et al., 2008) are sufficient to induce significant impairment of spatial working memory function.

There is evidence that persistent neural activity during delay periods of a working memory task is important for encoding the memory of a stimulus (reviewed in Constantinidis et al., 2018). When age-related alterations to the excitability properties of neurons comprising the working memory network occur, the network’s capacity for maintaining persistent activity—keeping activity tuned to the original stimulus—is impaired (Wang et al., 2011). To provide insight into the relationship between neuronal firing rate and synaptic changes seen in aging and working memory decline, we examined the parameter space of a bump attractor model of the DRT, then extended the model to simulate how aging might affect memory retention of multiple stimuli in a more complex task— the DRSTsp. The region of parameter space that simulated the DRT successfully was largest when the pyramidal neuron excitability parameter (*ν*_ce_) was tuned to fit the young FR data. The network model had its own built-in compensatory mechanisms: when pyramidal neuron excitability increased, the region with successful DRT models shifted to lower excitatory and higher inhibitory synapse weights. This tradeoff allowed networks to control the overall excitatory:inhibitory balance needed to maintain tuned persistent activity. These compensations were consistent with empirical data showing an age-related decrease in excitatory synapses, but not consistent with the reported decrease in inhibitory synapses (Peters et al., 2008). However, this compensatory mechanism led to lower firing rates during the delay period, as well as during the cue period when the stimulus current was also reduced, unifying our *in vitro* observations with the *in vivo* data (Wang et al., 2011). We predict that the network compensates higher FR in individual pyramidal neurons with aging by lower total synaptic input current to each neuron, and that the stimulus current could also be decreasing with aging.

To test how hyperexcitability of pyramidal neurons and loss of excitatory and inhibitory synapses with aging affected network output, we ran a second set of simulations: ‘aging’ model neurons by decreasing both excitatory and inhibitory synaptic strengths and increasing the FR of the excitatory neurons. DRT model performance (represented by the number of networks performing the task successfully) was substantially impaired in middle-aged and aged simulations, more than reported in a sample of rhesus monkeys (Wang et al., 2011), and the FR increase affected DRT performance much more than the loss of synapses did. Adding synaptic facilitation restored successful function to many middle aged and aged DRT models. However, only one of the aging cohort scenarios we tested captured the hyperexcitable pyramidal neuron *f*-I curves observed *in vitro*, dlPFC synapse loss observed with electron microscopy, and the *in vivo* FR decrease during the DRT cue and delay periods from Wang et al. (Wang et al., 2011): the one without synaptic facilitation but added hyperexcitability of *inhibitory* neurons to restore excitatory:inhibitory balance. One next step is to examine the excitability of dlPFC interneurons with aging, but our simulations raise several other questions about the DRT. Is there an age-related reduction in overall input to dlPFC during the cue period? How central a role does synaptic facilitation play during the task? How do age-related changes in synaptic plasticity, discussed below, affect DRT performance? Might neuromodulation (Arnsten et al., 2012; Davis et al., 2017; Wang et al., 2019) during working memory task performance reduce the pyramidal neuron hyperexcitability observed *in vitro*? Computational models provide an essential means for testing and refining hypotheses as future data become available.

Finally, we extended the DRT bump attractor model to create a model of working memory retention in a task like the DRSTsp. Other network models of working memory have simulated several stimuli at once (Edin et al., 2009; Wei et al., 2012; Mi et al., 2017). Yet, while the DRSTsp has been used for many years to evaluate spatial working memory capacity in rhesus monkeys (Herndon et al., 1997; Luebke et al., 2004; Chang et al., 2005; Moore et al., 2005; Peters et al., 2008; Luebke and Amatrudo, 2012; Moore et al., 2017)—including in the present study—to our knowledge this is the first computational model of this task. Increasing pyramidal neuron FR strengthened the response to the initial spatial cue and reduced the ability of the DRSTsp model networks to encode memories of subsequent spatial cues, so that middle-aged and aged networks had lower *DRST*_*sim*_ than young simulations. As with the DRT model, ‘aging’ the young DRSTsp model cohort led to severe impairment when increasing pyramidal neuron excitability alone, but not synapse loss alone. The maintenance of the DRSTsp model performance under synapse loss alone is consistent with Peters et al. (2008), which found no correlation between the numerical density of synapses and DRSTsp impairment. Simultaneously varying synapse loss and pyramidal neuron hyperexcitability within ranges observed with aging, both with and without hyperexcitability of interneurons, the amount of *DRST*_*sim*_ impairment was similar to that seen in DRSTsp scores across the adult rhesus monkey life span. Thus, our model predicts that pyramidal neuron hyperexcitability and synapse loss may be sufficient to explain empirically observed levels of spatial working memory impairment. Future experiments will examine whether the amount of synapse loss reported in (Peters et al., 2008) is also seen in the dlPFC of a subset of monkeys from this study, and whether there are correlations between those data, AP firing rates, and spontaneous postsynaptic currents.

There is an open debate about the role of synaptic mechanisms for working memory (Constantinidis et al., 2018; Lundqvist et al. 2018). While delay activity is recognized as important to working memory, a recent perspective posited that sparse spiking activity and synaptic plasticity between spike times—rather than asynchronous persistent activity—might be the mechanism actually responsible for memory maintenance (Lundqvist et al., 2018). This view is supported by several modeling studies (Sandberg et al., 2003; Mongillo et al., 2008; Lundqvist et al., 2011). Our models lend further support to an essential role for both persistent activity and synaptic facilitation for working memory tasks involving multiple cues. Synaptic plasticity (both facilitation and depression) help overcome limitations of continuous attractor models, such as filtering out distractors (Compte et al., 2000) and remembering multiple items (Itskov et al., 2011; Wei et al., 2012; Hansel and Mato, 2013; Rolls et al., 2013; Mi et al., 2017) as in our DRSTsp model. Bump attractor models of visuo-spatial working memory have shown that short-term synaptic facilitation in recurrent connections of excitatory neurons slows down the drift of the bump often seen in continuous attractor models (Itskov et al., 2011). Hansel and Mato (Hansel and Mato, 2013) showed that short-term synaptic facilitation might lead to the direction selectivity of synaptic weights and network bistability in a DRT model. *In vitro* experiments have shown synaptic facilitation in PFC pyramidal neurons (Hempel et al., 2000; Wang et al. 2006). Several other studies have shown age-related changes in synaptic plasticity (not modeled here), some correlating these changes with learning and memory deficits (Norris et al., 1998a, 1998b; Bach et al., 1999; Rosenzweig and Barnes, 2003; Thibault et al., 2007; Gant et al., 2011; Thibault et al., 2013; Gant et al., 2014; Gant et al., 2015).

This network-level study represents an extension of our past modeling studies at the single-neuron level on parameters affecting firing properties of individual pyramidal neurons with aging (Coskren et al., 2015; Luebke et al., 2015; Rumbell et al., 2016), in different cortical regions (Amatrudo et al., 2012), and in neurodegeneration (Goodliffe et al., 2018). The elegance of these network models lies in their ability to represent empirically observed changes in FR and synaptic weights with aging as perturbations to a small number of network parameters. Based on our findings with two *f*-I curves, replacing our firing rate model neurons with a spiking neuron model (Stein, 1967; Hodgkin and Huxley, 1990; Izhikevich, 2003; Teeter et al., 2018) in our DRT and DRSTsp networks should yield qualitatively similar results. Our future network modeling will incorporate age-related changes in passive and active channel parameters predicted in our and other past studies (Wang et al., 2011; Coskren et al., 2015; Rumbell et al., 2016), and other empirically observed changes in aging dlPFC including increased slow afterhyperpolarization current (Luebke and Amatrudo, 2012); white matter changes (review: Luebke et al., 2010; Kubicki et al., 2019); and dysregulation of calcium homeostasis (Foster, 2007; Toescu and Vreugdenhil, 2010; Oliveira and Bading, 2011). Age-related changes to synaptic strength and neurotransmitters reduced the probability of short-term memory recall from a discrete attractor network model (Rolls and Deco, 2015); similar perturbations in a spiking-network version of our DRT and DRSTsp models would be extremely insightful.

The main limitation of our current DRSTsp model—assuming that the network correctly identifies the novel stimulus among several retained—is also a limitation of the classical bump attractor as a whole. Experiments have shown that decision-making and attentional tasks involve connections between dlPFC and other cortical regions including posterior parietal cortex, anterior cingulate cortex, and secondary motor areas (Medalla and Barbas, 2010; Oemisch et al., 2015; Constantinidis and Qi 2018; Marvel et al. 2019; Tsunada et al. 2019). These connections may represent different aspects of a task like DRSTsp, such as the monkey’s internal model of itself and the task; its level of attention; signals of reward and error; and motor traces that might correspond to rehearsal strategies between active stimulus stages of the DRSTsp. Recent theoretical studies have shown that model neurons with mixed selectivity, responding to sensory stimuli as well as internal states of the system, can perform complex cognitive tasks (Maass et al., 2002; Salinas, 2004; Rigotti et al., 2010; Chaisangmongkon et al., 2017). Such units could be well-suited for extending the DRSTsp model introduced here to simulate choice of the novel spatial cue and related internal signals, plus the full 3×6 array of wells presented to the monkeys during the DRSTsp. There is evidence that aging affects information flow between cortical regions, which can contribute to working memory impairment (Engle et al., 2016; Lee et al., 2016; Proskovec et al., 2016; King et al., 2018; Koen et al., 2019). Any such age effects must be examined in concert with the changes to dlPFC synapses and neuronal excitability, to predict the extent to which these concomitant changes compound versus partially compensate each other.

In summary, our computational models of two spatial memory tasks predicted that empirically observed changes in pyramidal neuron excitability and synapse loss lead to spatial working memory impairment with aging. Computational studies such as this provide a means to evaluate connections between physiological and behavioral—and *in vitro* vs. *in vivo*—experiments, extending the impact provided by animal studies.

## AUTHOR CONTRIBUTION

SI developed and implemented the models, analyzed the results, carried out the statistical analysis of the data and wrote the manuscript. JIL performed the experiments, wrote the manuscript and designed the study. WC performed the experiments. DD designed the statistical analysis of the data. CMW carried out the statistical analysis of the data, wrote the paper and designed the study.

## FUNDING

This work was supported by NIH/NIA grants R01 AG059028 and R56 AG049870.

## ACKNOWLEDGEMENTS

We thank Drs. Tara Moore and Doug Rosene for the behavioral data and brain tissue from which recordings were made. We thank Drs. Maya Medalla, Patrick Hof, and Patrick Coskren for productive discussions.

