## Supplementary Material for "Network models predict that pyramidal neuron hyperexcitability and synapse loss in the dlPFC lead to age-related spatial working memory impairment in rhesus monkeys"

### 1. DRT MODEL SIMULATIONS WITH AN ALTERNATIVE *F*-I CURVE

We compared LHS results for an alternative, continuous activation function that satisfies the physiological requirements (see Brunel 2003) but is less excitable:

$$f_g(I) = \begin{cases} 0 & I \leq 0 \\ \nu_c \left(\frac{I}{I_c}\right)^p & 0 < I < I_c \\ p \cdot \nu_c \left(\frac{I}{I_c} - \frac{p^p - 1}{p^p}\right)^{1/p} & I_c \leq I \end{cases}.$$

This function generalizes the suprathreshold regime (when $I_c \leq I$), still with a concave shape and

$$p \cdot \nu_c \left(\frac{I}{I_c} - \frac{p^p - 1}{p^p}\right)^{1/p} = \nu_c$$

when $I = I_c$, but given by a root function of general order instead of a square root.

The DRT model simulations were repeated using $f_g(I)$ for the LHS regime in Table 2 in the main document (Supp. Fig. 1). We found good fits for the empirical data of the young, middle-aged and aged groups of subjects for $\nu_{ce}$ = 2.63 Hz, $I_{ce}$ = 122 pA, and $p_e$ = 4.9, 6.14, and 7.11, respectively (Supp. Fig. 1a). Therefore, with this activation function, varying $p_e$ was sufficient to fit the empirical *f*-I curves of the three age groups. Setting $\nu_{ci}$ = 2.63 Hz, $I_{ci}$ = 118 pA, and $p_i$ = 50 for the network inhibitory neurons, we repeated the DRT simulations at each of the 4200 points in the LHS for $p_e$ = 3, 4, 5, 6, 7, and 8. As when $\nu_{ce}$ increased in the classical (but more excitable) *f*-I curve in the main text, the number of points maintaining TPA-S decreased as $p_e$ increased. Overall, points sampled in the regime became more excitable: Supplemental Figure 1d shows fewer under-excited and more over-excited cases as $p_e$ increased (blue vs. red dots). Most synaptic weight trends reported in the main text were also maintained here, suggesting that our results are robust to variations in the f-I curve formulation.

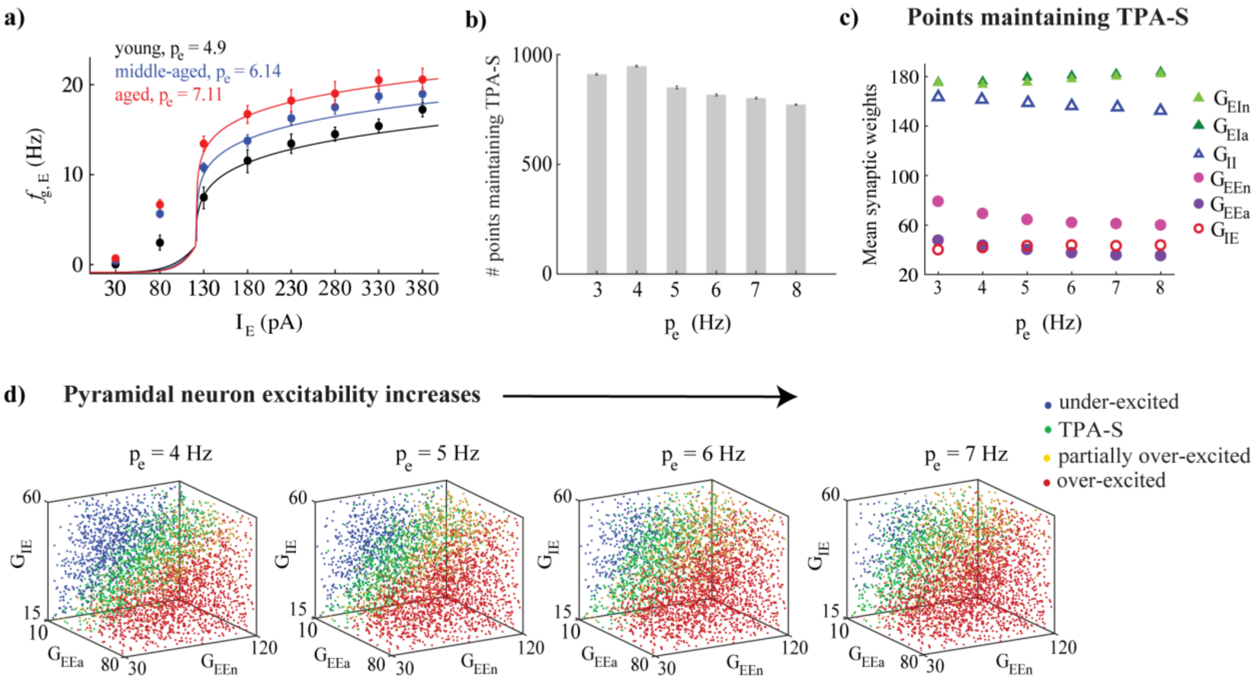


**Supplementary figure 1: DRT model output with $f_g(I)$ varied across the parameter space.** a) $f_g(I)$ fit (solid lines) to empirical AP firing rates of pyramidal neurons of young, middle-aged and aged subjects, averaged for each age group (black, blue, and red respectively, shown as mean ± SEM at each injection level). b) Number of points in each LHS maintaining TPA-S as $p_e$ increased. c) Mean synaptic weights ($G_{EEa}$, $G_{EEn}$, $G_{IE}$, $G_{EIa}$, $G_{EIn}$, and $G_{II}$) for all points maintaining TPA-S as $p_e$ increased. $G_{EEa}$, $G_{EEn}$, and $G_{IE}$ values shown as purple, pink, and red open circles respectively; $G_{EIa}$, $G_{EIn}$, and $G_{II}$ values shown as dark green, light green and open triangles respectively. S.E.M. bars lie beneath the symbols. d) DRT model output for 4200 points of the parameter space LHS, shown in 3D projections across the subspace of the excitatory ($G_{EEa}$ and $G_{EEn}$) and the inhibitory ($G_{IE}$) synaptic weights of pyramidal neurons as their excitability increased ($p_e$ = 4, 5, 6, and 7). Blue, green, yellow, and red dots represent under-excited networks, networks maintaining TPA-S, and partially and completely over-excited networks respectively.
